# The analysis of the role of MexAB-OprM on quorum sensing homeostasis shows that the apparent redundancy of Pseudomonas *aeruginosa* multidrug efflux pumps allows keeping the robustness and the plasticity of this intercellular signaling network

**DOI:** 10.1101/2020.03.10.986737

**Authors:** Manuel Alcalde-Rico, Jorge Olivares-Pacheco, Nigel Halliday, Miguel Cámara, José Luis Martínez

**Author notes:** Corresponding authors: JO, JLM.

## Abstract

Multidrug efflux pumps are key determinants for antibiotic resistance. Besides contributing to intrinsic resistance, their overexpression is frequently a cause of the increased resistance acquired during therapy. In addition to their role in resistance to antimicrobials, efflux pumps are ancient and conserved elements with relevant roles in different aspects of the bacterial physiology. It is then conceivable that their overexpression might cause a burden that will be translated into a fitness cost associated with the acquisition of resistance. In the case of *Pseudomonas aeruginosa*, it has been stated that overexpression of different efflux pumps is linked to the impairment of the quorum sensing (QS) response. Nevertheless, the causes of such impairment are different for each analyzed efflux pump. In this study, we performed an in-depth analysis of the QS-mediated response of a *P. aeruginosa* antibiotic resistant mutant that overexpresses MexAB-OprM. Although previous work claimed that this efflux pump extrudes the QS signal 3-oxo-C12-HSL, we show otherwise. Our results suggest that the observed attenuation in the QS response when overexpressing this pump is related to a reduced availability of intracellular octanoate, one of the precursors of the biosynthesis of alkyl quinolone QS signals. The overexpression of other *P. aeruginosa* efflux pumps has been shown to also cause a reduction in intracellular levels of QS signals or their precursors impacting on these signaling mechanisms. However, the molecules involved are distinct for each efflux pump, indicating that they can differentially contribute to the *P. aeruginosa* quorum sensing homeostasis.

**Importance:** The success of bacterial pathogens to cause disease relies on their virulence capabilities as well as in their resistance to antibiotic interventions. In the case of the important nosocomial pathogen *Pseudomonas aeruginosa*, multidrug efflux pumps participate in the resistance/virulence crosstalk since, besides contributing to antibiotic resistance, they can also modulate the quorum sensing (QS) response. We show that mutants overexpressing the MexAB-OprM efflux pump, present an impaired QS response due to the reduced availability of the QS signal precursor octanoate, not because they extrude, as previously stated, the QS signal 3-oxo-C12-HSL. Together with previous studies, this indicates that, although the consequences of overexpressing efflux pumps are similar (impaired QS response), the mechanisms are different. This ‘apparent redundancy’ of RND efflux systems can be understood as a *P. aeruginosa* strategy to keep the robustness of the QS regulatory network and modulate its output in response to different signals.

## Introduction

*Pseudomonas aeruginosa* is an opportunistic pathogen of special concern due to its capability to produce a large variety of serious human infections and to its low susceptibility to different antibiotics [1, 2]. It is worth noticing that genes that contribute to *P. aeruginosa* intrinsic antibiotic resistance are, in several occasions, key components of bacterial physiology [3, 4]. This is the case of the Resistance, Nodulation and cell-Division (RND) family of efflux pumps, which, besides being important mechanisms of antibiotic resistance in *P. aeruginosa* [5, 6], also may play a key role in its pathogenesis and adaption to host environment [3, 7]. In this work, we address the role of the MexAB-OprM efflux system, one of the most relevant RND systems for intrinsic and acquired antibiotic resistance of *P. aeruginosa* [8, 9], on the modulation of quorum sensing (QS) responses in this bacterium.

The QS response consists in a population-scale cooperative behavior promoted by cell-to-cell communication systems that control the expression of a large set of genes in a cell-density way [10]. This intercellular communication system is based on the synthesis, delivery and progressive accumulation of autoinducer compounds, known as QS signal molecules (QSSMs), which are recognized by specific cell receptors. When QSSMs concentrations reach a threshold, the QS response is triggered in the population. This population-scale response regulates a wide number of diverse physiological processes [11], including production of private and public goods [12], biofilm formation [13], host-bacteria interactions [14] and virulence factors production [15]. The QS responses usually impose a fitness burden at single cell level but with social benefit at population level [16, 17].

The QS regulatory network of *P. aeruginosa* is based on the production of two different kinds of QSSMs: the *N*-acyl-L-homoserine lactones (AHLs) and the 2-alkyl-4(1*H*)-quinolones (AQs) [10]. These signals are synthesized and detected by the Las, Rhl and Pqs regulation systems. The Las system is based on the production of *N*-(3-oxododecanoyl)-L-homoserine lactone (3-oxo-C12-HSL) by LasI synthase and the detection of this signal by the LasR transcriptional regulator. The *N*-butanoyl-L-homoserine lactone (C4-HSL) is part of the Rhl system being synthesized by RhlI and sensed by the RhlR regulator. The synthesis of the Pseudomonas Quinolone Signal (PQS), and its immediate precursor, 2-heptyl-4-hydroxyquinoline (HHQ), requires the enzymes encoded by the *pqsABCDE* operon and *pqsH*. These two QSSMs are detected by the LysR-type transcriptional regulator PqsR (also known as MvfR) and are the main QSSMs associated with the Pqs QS system. The interconnection between these QS systems has been described as hierarchized, with the Las system at the top controlling the activity of the Rhl and Pqs systems, which then modulate both their own activity and the expression of the other elements of the QS system [10]. However, there are studies suggesting that the hierarchy can be affected by other regulators, growth conditions or through the activity of other elements not directly involved in the canonical QS-regulatory network [18–22].

Further, there are key elements like PqsE that may function as QS modulators without binding to any QSSM, giving a higher level of complexity to the QS regulatory network [20, 21, 23, 24]. Another category of modulators of the QS response is formed by multidrug efflux pumps. Indeed, previous works have reported that the RND efflux systems MexAB-OprM, MexCD-OprJ, MexEF-OprN and MexGHI-OpmD are involved in the modulation of the *P. aeruginosa* QS response [25–29]. In the case of MexAB-OprM, whose over-expression leads to a lower production of some QS-regulated phenotypes [25, 30], it has been described that this efflux pump is able to extrude 3-oxo-C12-HSL [26]. However, our results show that overproduction of MexAB-OprM impairs the QS-mediated response due to the reduction in the availability of the AQs biosynthetic precursor octanoic acid [31] and consequently a reduced AQs production, and not to a high non-physiological extrusion of 3-oxo-C12-HSL

Therefore, the results of the present study, together with previously published works, evidence that the underlying causes of the impaired QS response and the lower production of QSSMs observed in MexAB-OprM, MexCD-OprJ and MexEF-OprN overexpressing mutants are different. We propose that the “apparent redundancy” observed between these RND systems in their modulation of *P. aeruginosa* QS response is not casual and could be employed by *P. aeruginosa* to fine tuning its QS regulatory network in response to changes in environmental conditions and nutritional requirements.

## Materials and methods

### Bacterial strains, plasmids and primers

All the bacterial strains and plasmids used in this work are shown in Table 1. Primers used are listed in Table 2.

**Table 1.**
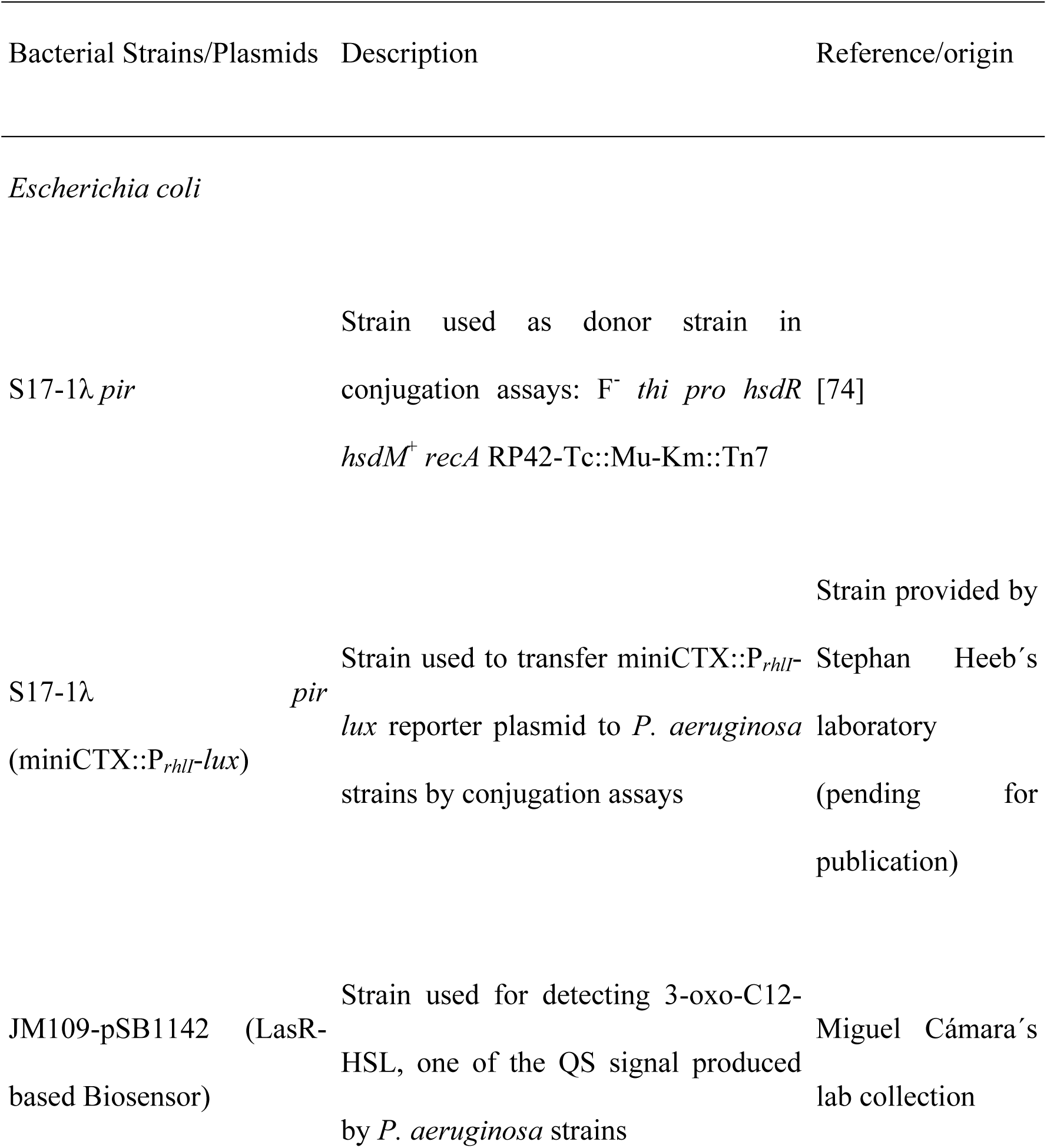
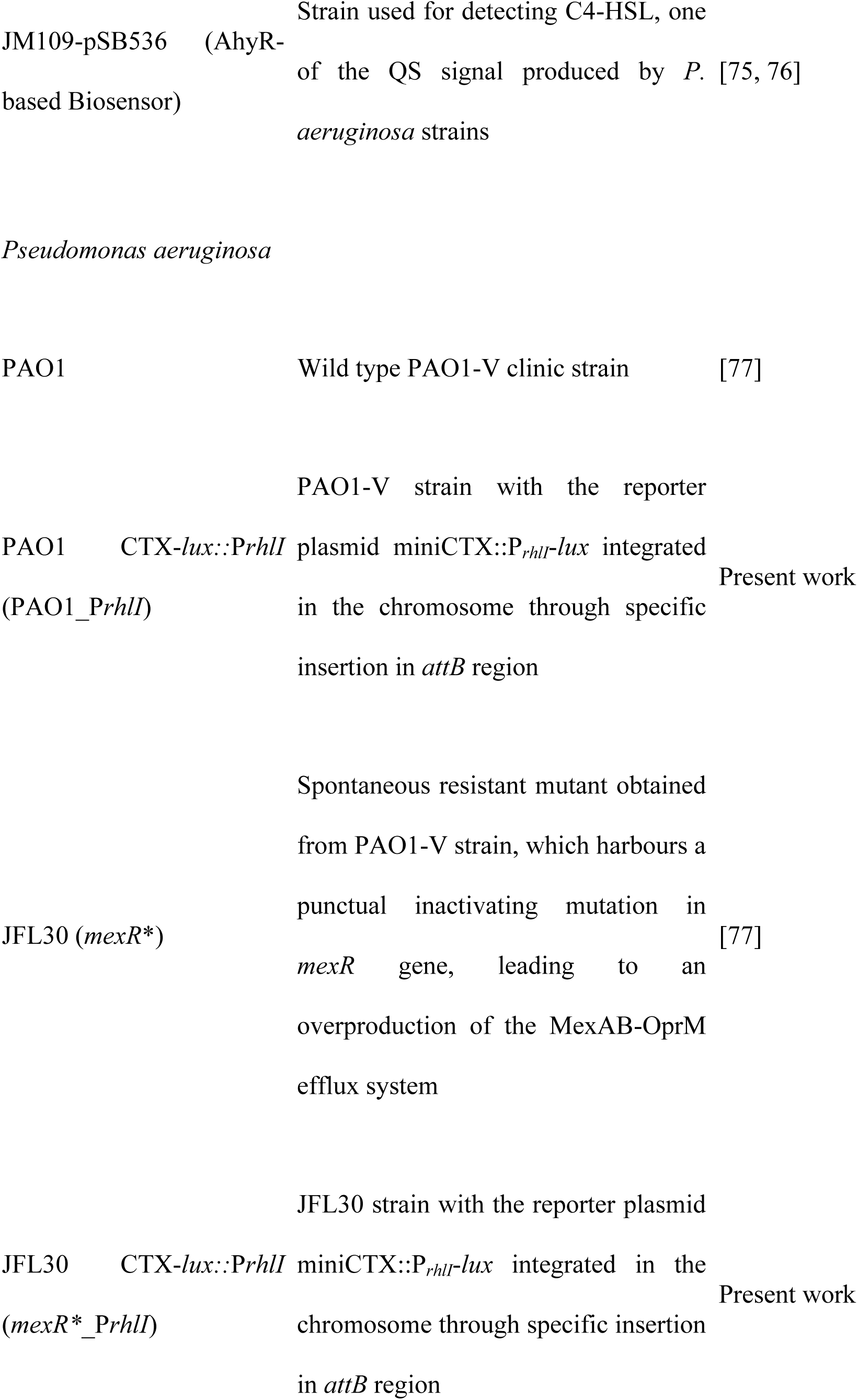
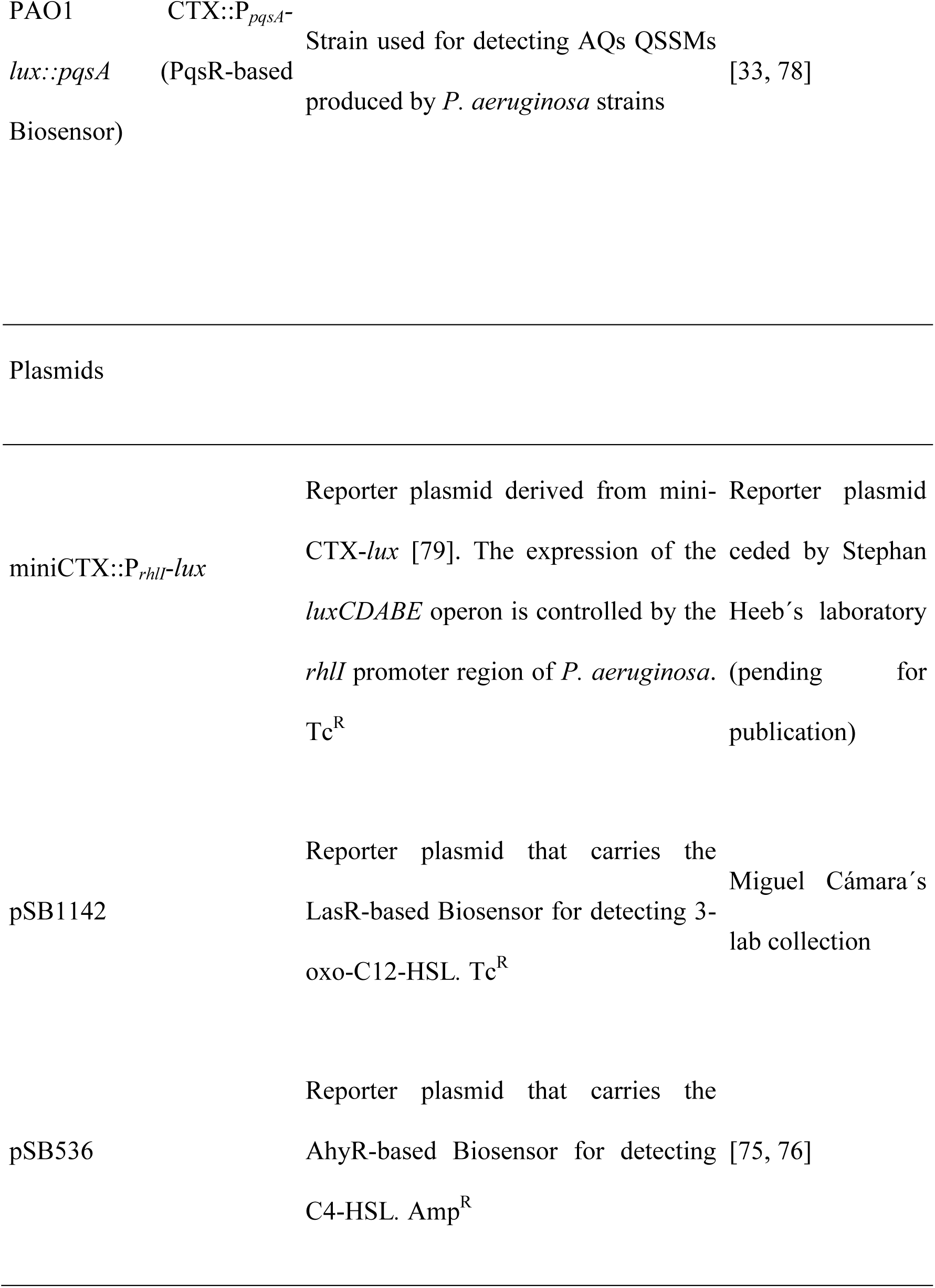
Bacterial strains and plasmids used in the present work.

**Table 2.**
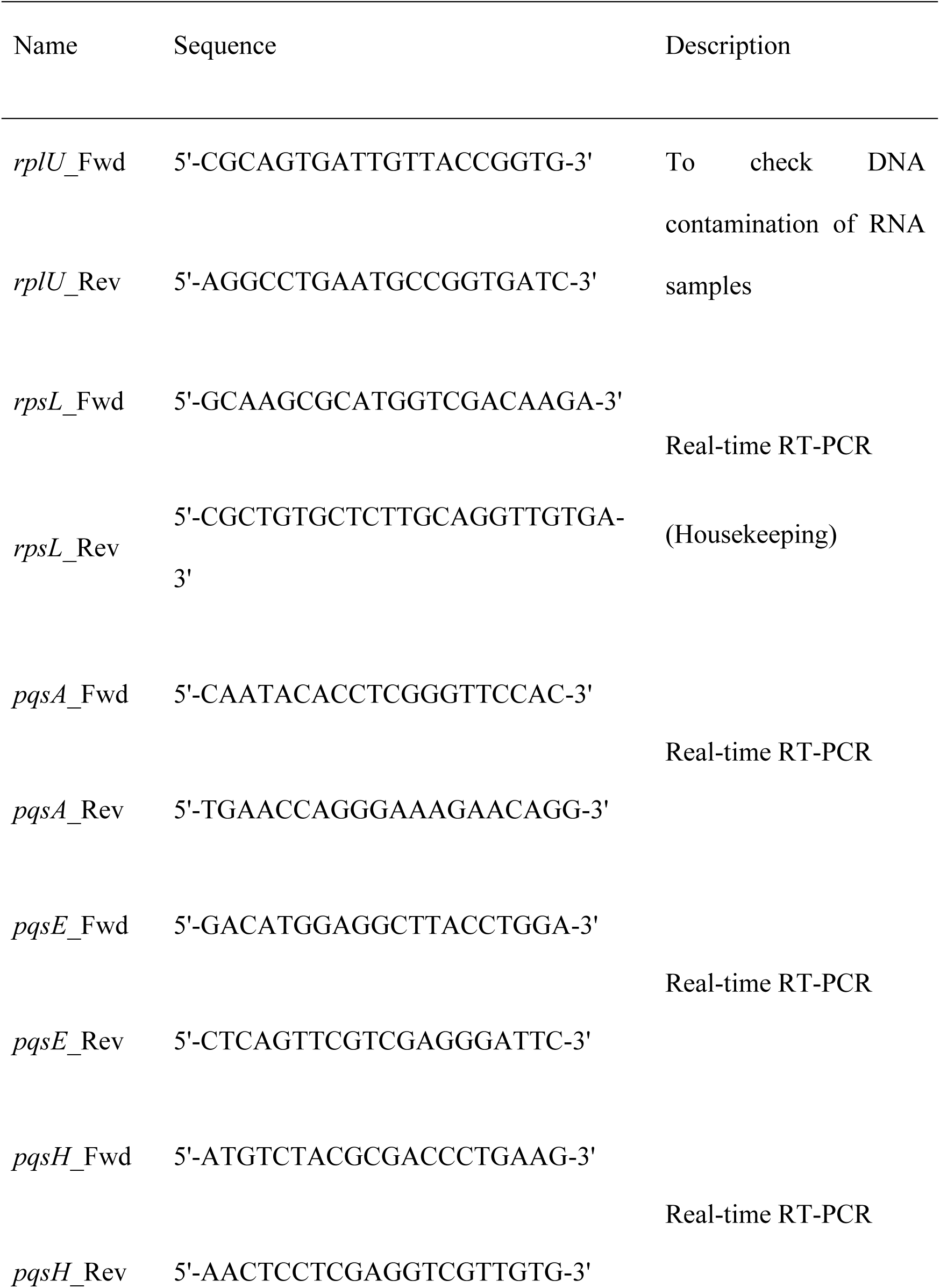
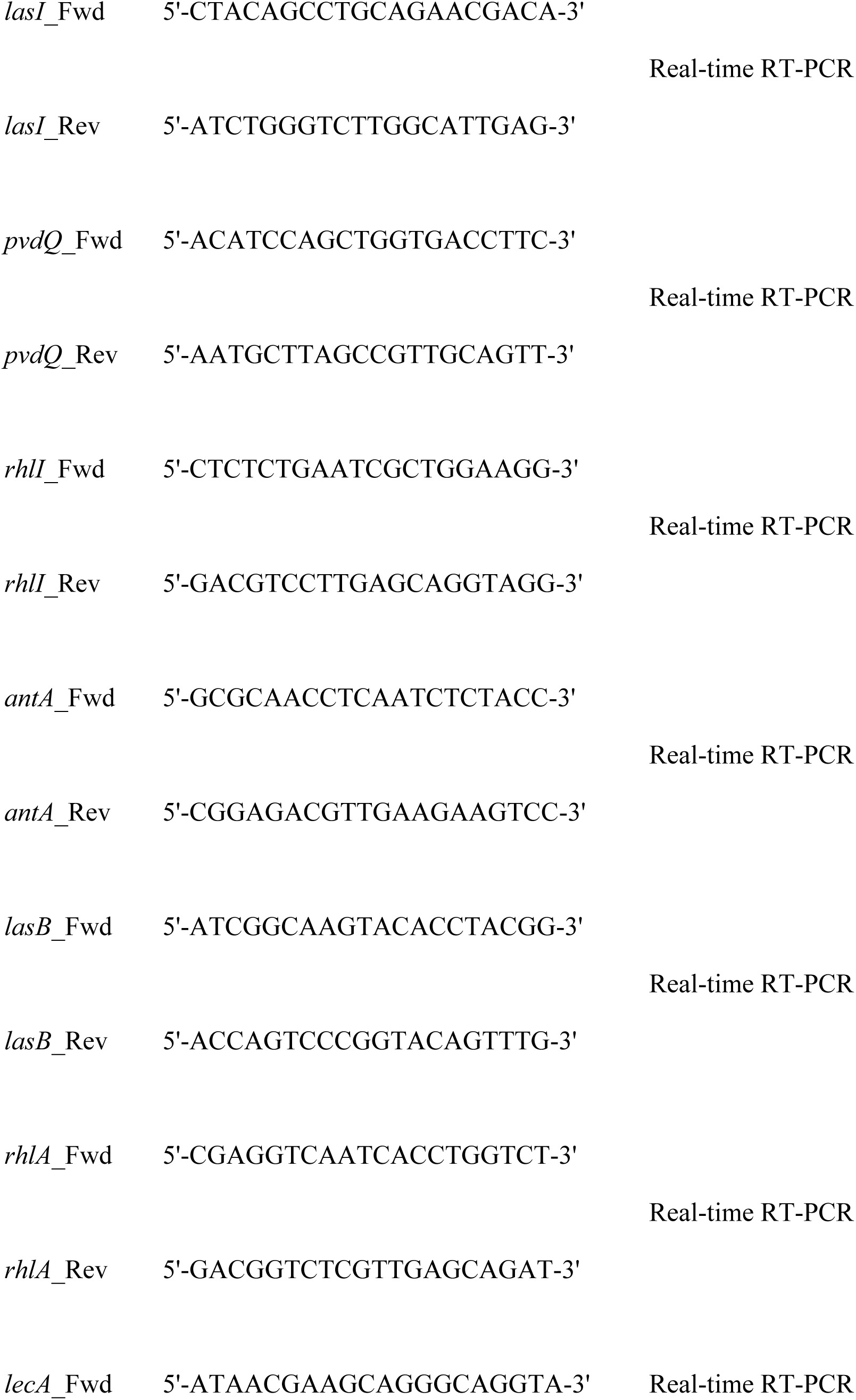
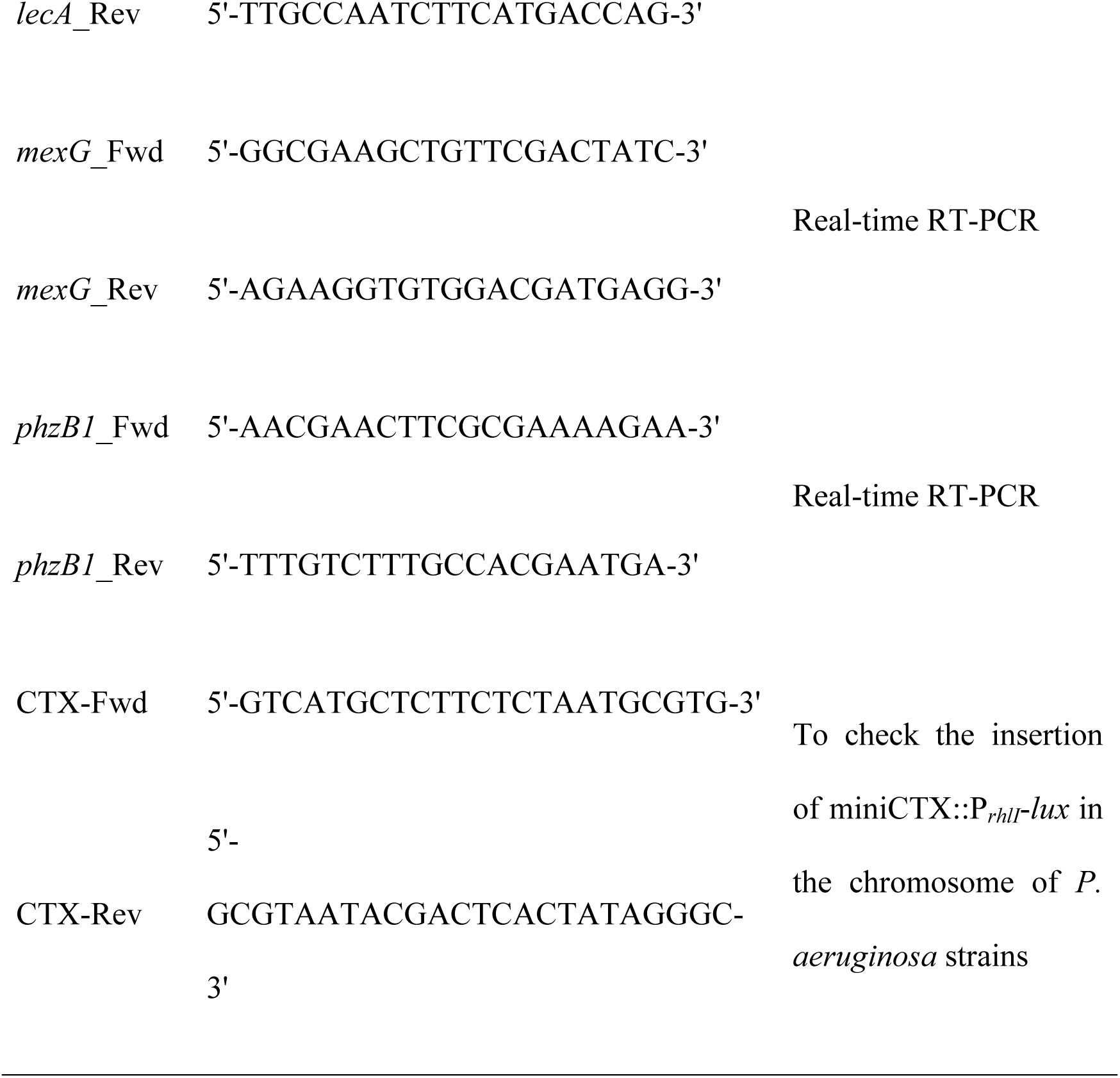
Primers used in the present work.

### Growth media and culture conditions

The experiments were carried out in 100 ml glass-flasks containing 25 ml of broth, which were inoculated at the beginning of the experiments at OD_600_ = 0.01, unless other conditions are stated. All strains were routinely cultured in Lysogeny Broth (LB) Lennox (Pronadisa) at 37 °C with shaking (250 rpm). For the growth of *E. coli* strains containing plasmids with an ampicillin (Amp) resistance marker, LB medium containing Amp 100 μg/ml was used. For the growth of *E. coli* or *P. aeruginosa* strains harboring the miniCTX::P*_rhlI_*-*lux* constructions, growth media containing tetracycline (Tc) 10 μg/ml or 100 μg/ml were used, respectively. LB medium containing anthranilate 1 mM (Acros organics, Thermo Fisher Scientific) was obtained using a stock solution of anthranilate 100 mM adjusted to pH = 7.2. For the supplementation of the LB medium with octanoate 5 mM, the corresponding amount of sodium octanoate (Sigma, Aldrich) was directly dissolved in LB medium and sterilized by filtration through a membrane with a 0.22 μm pore size. For detecting of the QSSMs by Thin Layer Chromatography (TLC) (see below), two different semi-solid media were used according to Yates *et al.* (2002) [32] and Fletcher *et al.* (2007) [33].

### Pyocyanin extraction

The *P. aeruginosa* cultures were incubated along 20 hours at 37 °C with shaking and the accumulation of pyocyanin in cell-free supernatants was determined following the method described by Essar et al. [34]. The concentration of pyocyanin was calculated based on its molar extinction coefficient.

### Total RNA extraction, sequencing and analysis of transcriptomes

Overnight cultures of *P. aeruginosa* were washed and diluted in LB to an OD_600_ of 0.01. They were incubated up to exponential phase of growth (OD_600_ = 0.6) and then were diluted again to an OD_600_ of 0.01. The cultures were grown to reach the exponential (OD_600_ = 0.6) or early stationary phase of growth (OD_600_ = 2.5), time points in which total RNA was obtained as described Schuster *et al*. (2003) [35] using the RNeasy mini kit (QIAGEN). Before RNA extraction, the samples for RNA sequencing were treated with “RNAProtect Bacteria Reagent” (QIAGEN) following the manufacturer’s instructions. For Real-Time reverse transcription PCR (RT-PCR) assays, the RNA extractions were carried out in triplicate. To discard possible DNA contamination, a PCR reaction using the PCR Master Mix (Promega) and the primers *rplU*_Fwd and *rplU*_Rev (Table 2) was carried out. RNA samples were sequenced at the “Centro Nacional de Análisis Genómico” (CNAG), Barcelona (Spain). Ribosomal RNA was removed using “RiboZero rRNA Removal kit for Bacteria”. To generate the libraries, 2 μg of RNA were treated with “TruSeq RNA sample preparation kit” (Illumina) combined with a specific strand labeling using dUTPs [36]. The sequencing in 2 x 75 pair-end format with Illumina technology was performed. The sequences were aligned against the PAO1 reference genome NC_002516 available in the “Pseudomonas Genome Database” (PseudoCAP) [37] and gene expression was quantified using the “CLC Genomics Workbench” software (QIAGEN). The numeric value of gene expression was normalized to Reads Per Kilobase of gene per million Mapped reads (RPKM). Subsequently, a cut-off value of 1 was added to each RPKM (RPKM + 1) in order to minimize the misleading fold change values caused by RPKMs close to 0 [38]. The LogRatio parameter of each gene was calculated using the formula LogRatio = Log_2_ (RPKM*_mexR*_*/RPKM_PAO1_). Genes with LogRatio values higher than 1 or lower than −1 were considered to be affected in their expression and were grouped in the functional classes established in PseudoCAP [37].

### Real-time Reverse-Transcription Polymerase Chain Reaction (RT-PCR)

cDNA synthesis was carried out using the High Capacity cDNA reverse transcription kit (Applied Biosystems). The RT-PCR reactions were performed in an ABI prism 7500 system” (Applied Biosystems) using the Power SYBR green kit (Applied Biosystems) following manufacturer’s instructions, and adding to each reaction 50 ng of cDNA and 400 nM of paired-primers (Table 2). The housekeeping gene *rpsL* was used for normalization and gene expression was quantified using the 2^−ΔΔCt^ method [39].

### LC-MS/MS Analysis of QSSMs in Supernatants

The extraction of QSSMs from supernatants of *P. aeruginosa* cultures grown in LB to reach the late stationary phase (16 hours of incubation) and their identification and quantification by LC-MS/MS was performed as previously described [40]. In essence, the efficiency and reproducibility of QSSMs extraction were monitored by the addition of a deuterated AHL-internal standard, d9-C5-HSL, and a deuterated AQs-internal standard, d4-PQS, to each sample prior to the extraction protocol. The analyte peak areas obtained for each one of the QSSMs analyzed in a specific sample were normalized with respect to those obtained for the corresponding internal standard in the same sample.

### Thin Layer Chromatography (TLC) and Time-Course Monitoring (TCM) of QSSMs accumulation

The extraction of QSSMs was carried out following the methodology detailed by Alcalde-Rico *et al.* (2018) [27]. The TLC-based detection of AHLs and AQs were performed following the instructions described by Yates *et al.* (2002) [32] and Fletcher *et al.* (2007) [33], respectively. The TCM-based detection of QSSMs were carried out following the methodology described by Alcalde-Rico *et al.* (2018) [27]. For TLC-spots quantifications by densitometry, three independent measurement for each sample and TLC were obtained using the “ImageJ” software.

### Site-specific insertion of the miniCTX::P*_rhlI_*-*lux* reporter in the chromosome of *P. aeruginosa* strains and real-time analysis of P*rhlI* activation

The transfer of the miniCTX::P*_rhlI_*-*lux* plasmid and its integration into the chromosome of *P. aeruginosa* was carried out following the methodology described by Hoang *et al.* (2000) [41]. Overnight cultures of donor (*E. coli* S17-1λ *pir* (miniCTX::P*_rhlI_*-*lux*)) and recipient strains (*P. aeruginosa* PAO1 and *mexR*\*) were washed with LB medium and mixed prior to be poured over LB plates, which were incubated for 8 hours. *P. aeruginosa* transconjugants were selected using Petri dishes with *Pseudomonas* Isolation Agar (PIA, Fluka) containing Tc 100 μg/ml. The chromosomal insertion of miniCTX::P*_rhlI_*-*lux* was checked by PCR using the CTX-Fwd and CTX-Rev primers (Table 2). The analysis of the P*rhlI* promoter activity was carried out in Flat white 96-well plates with clear bottom (Thermo Scientific Nunc) and each reporter strain was inoculated to an OD_600_ of 0.01 in 200 µl of LB. The luminescence and OD_600_ were monitored every 10 minutes for 20 hours using a multi-plate reader (TECAN infinite 200). The mean values correspond to the average of three biological replicates and the error bars represent their standard deviation.

### Statistical Analysis

The area under the curve of each biological replicate was quantified using the GraphPad Prim software. All the experiments, in which Student’s two-tailed test with a confidence interval of 95% were applied to analyze statistical significance, were performed at least in triplicate. Variations with a P-value lower than < 0.05 were considered significant (* represent P-values < 0.05; ** P-values < 0.01; *** P-values < 0.001).

## Results

### Effect of MexAB-OprM overexpression on the transcriptome of *P. aeruginosa*

To analyze the global effect of MexAB-OprM overexpression on *P. aeruginosa* physiology, the transcriptomes of both wild-type PAO1 strain and a *mexR*\* mutant, which harbors an inactivating mutation in the *mexAB-oprM* repressor gene, *mexR*, were analyzed by RNAseq and compared. Since it has been described that MexAB-OprM is involved in the QS response [25, 26, 42] and its expression is growth phase regulated [42, 43], the transcriptomic assays were addressed in exponential (OD_600_ = 0.6) and early stationary phase of growth (OD_600_ = 2.5).

The overexpression of MexAB-OprM in the *mexR*\* mutant is associated with significant changes in the expression of 182 and 346 genes during exponential and stationary growth phase, respectively (Table S1). The genes affected were grouped based on the functional classes assigned in PseudoCAP [37]. The major functional class affected by MexAB-OprM overexpression in both growth phases was the one corresponding to secreted factors, whose genes were mainly expressed at a lower level in the MexAB-OprM overexpressing *mexR\** mutant than in the wild-type PAO1 strain (Figure 1). Other functional classes strongly affected in both phases of growth were: i) the biosynthesis of prosthetic groups, cofactors and carriers; ii) antibiotic resistance and susceptibility; iii) those involved in adaptation and protection; iv) genes encoding enzymes involved in carbon metabolism; and v) genes involved in the metabolism of fatty acids and phospholipids. Many of genes, whose expression was altered in the *mexR\** mutant in both phases of growth, belong to the QS regulatory network. Among them, key components in AQs biosynthesis such as *pqsA*, *pqsB*, *pqsD*, *pqsE*, *phnA* and *phnB*, as well as genes regulated by Pqs system such as *phzA2*, *phzB1*, *rhlA*, *rhlB* or *lasB* were found (Table 3).

**Figure 1.**
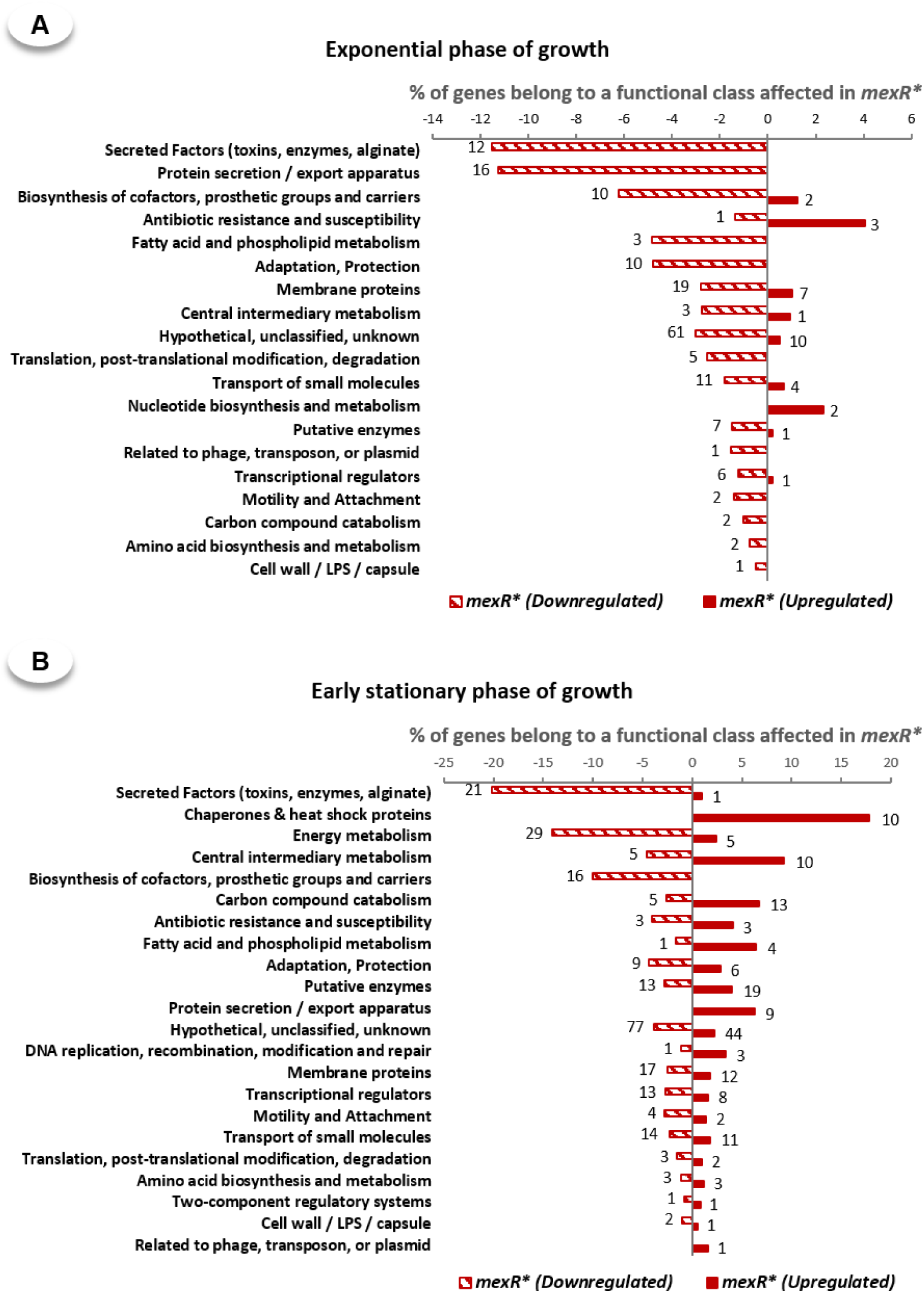
Genes affected in the MexAB-OprM overproducer mutant along both the exponential and early stationary phase of growth. The total RNA was extracted from PAO1 and *mexR*\* cultures at both (A) exponential (OD_600_ = 0.6) or (B) early stationary phase of growth (OD_600_ = 2.5) as is described in Methods. The expression value for each gene was calculated based on its RPKM (Reads Per Kilobase of gene per million Mapped reads) and only RPKM changes over or below two-fold with respect the control condition were considered as significant variation of the gene expression. The genes affected were grouped in the corresponding functional class stablished in PseudoCAP and the number of the genes affected in each functional class is represented near to the corresponding bar. The percentage of genes whose expression is affected in each functional class was calculated over the total genes grouped in the same category. The results showed that overexpression of the MexAB-OprM system has a strong impact over the transcriptome of *P. aeruginosa* in the both growth phases analyzed and that the most affected functional class is “Secreted Factors (toxins, enzymes, alginate)”.

**Table 3.**
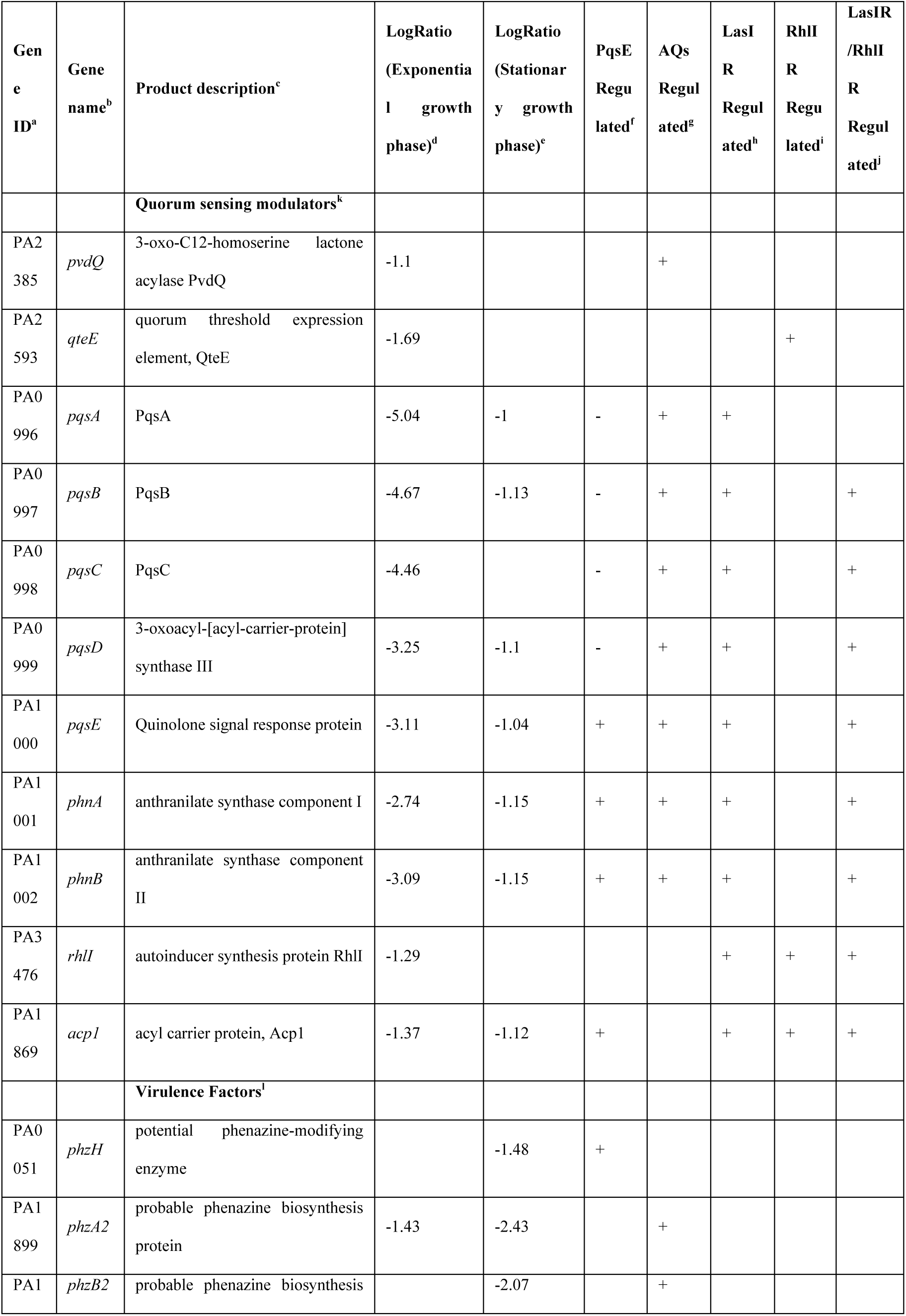
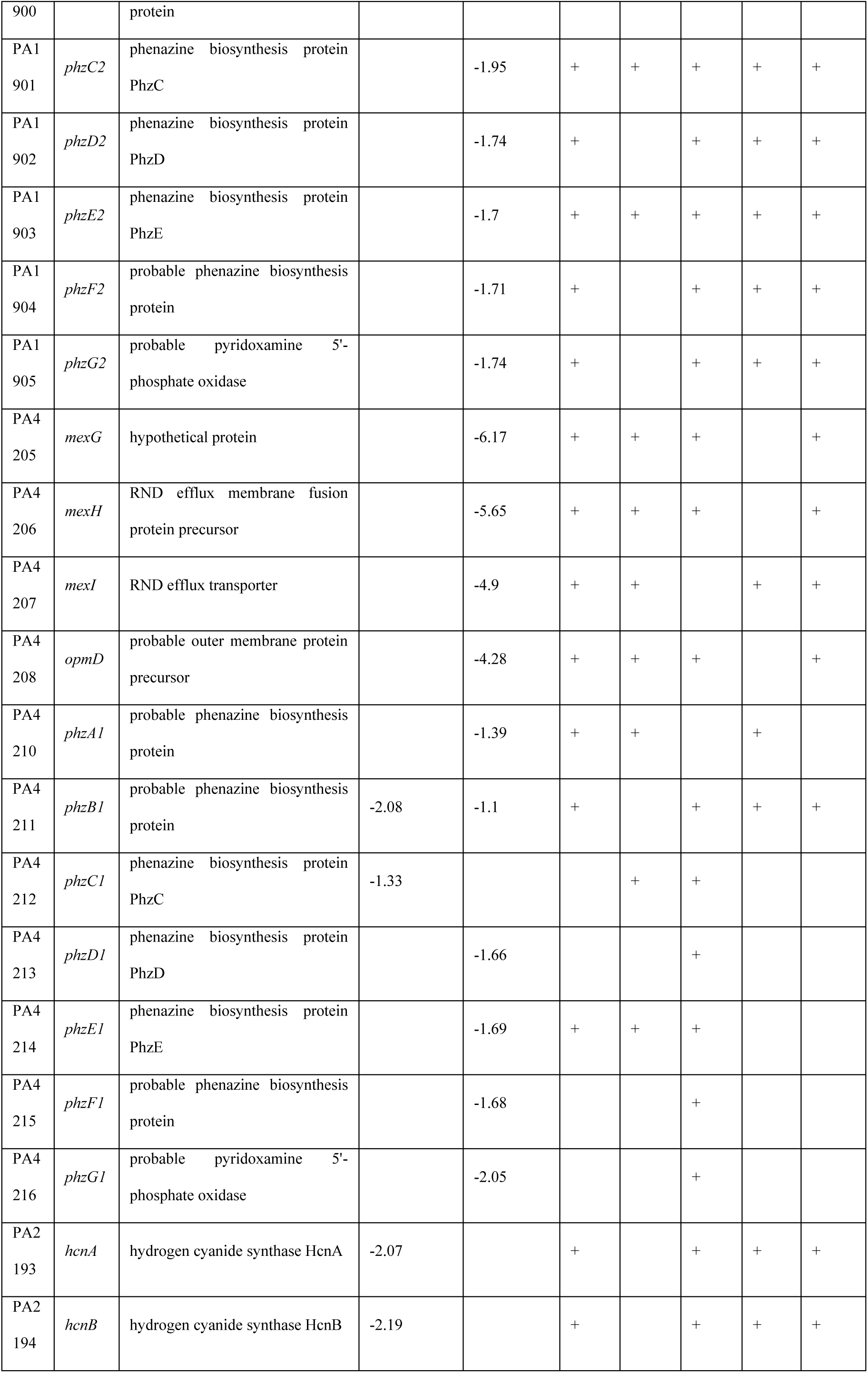
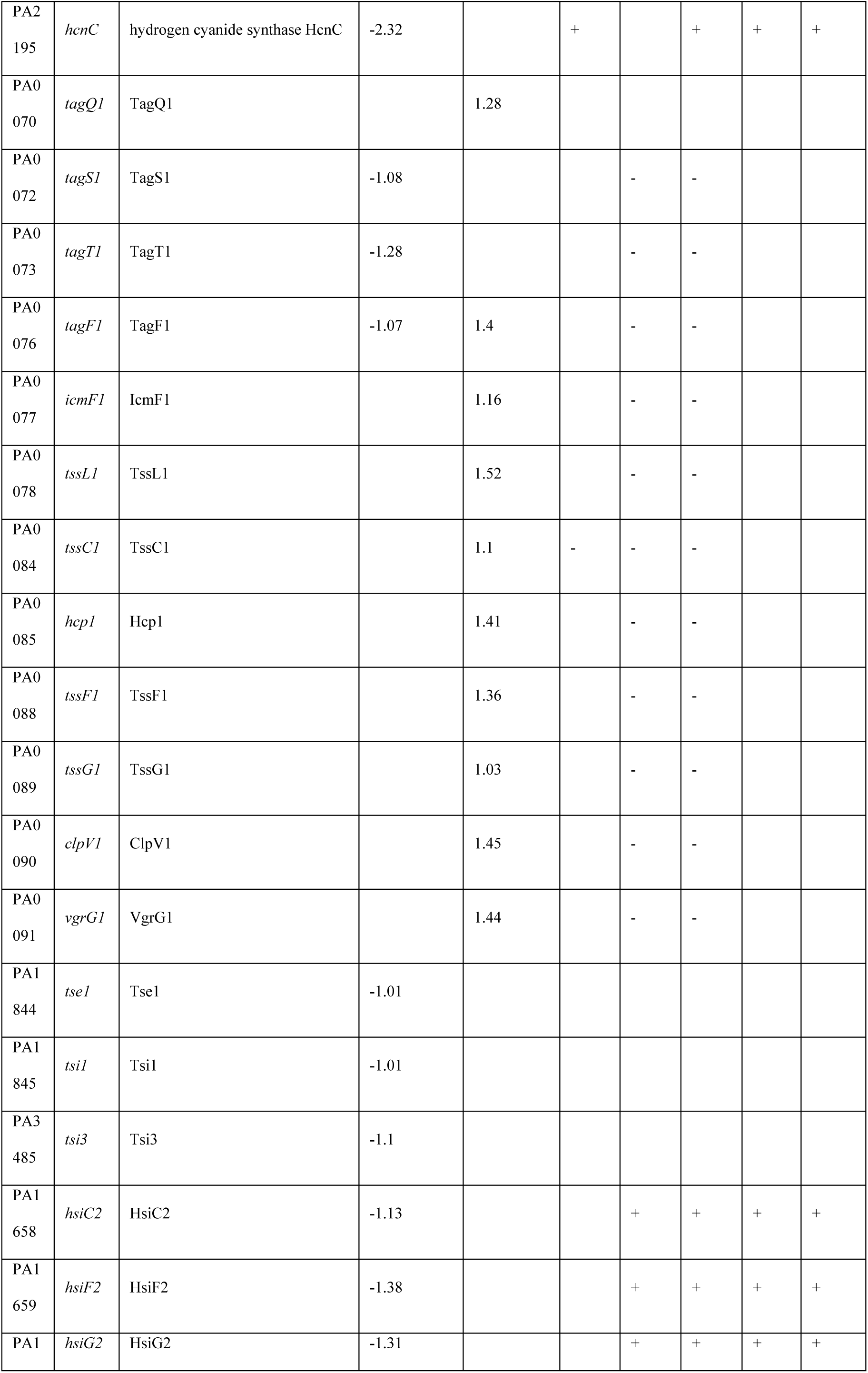
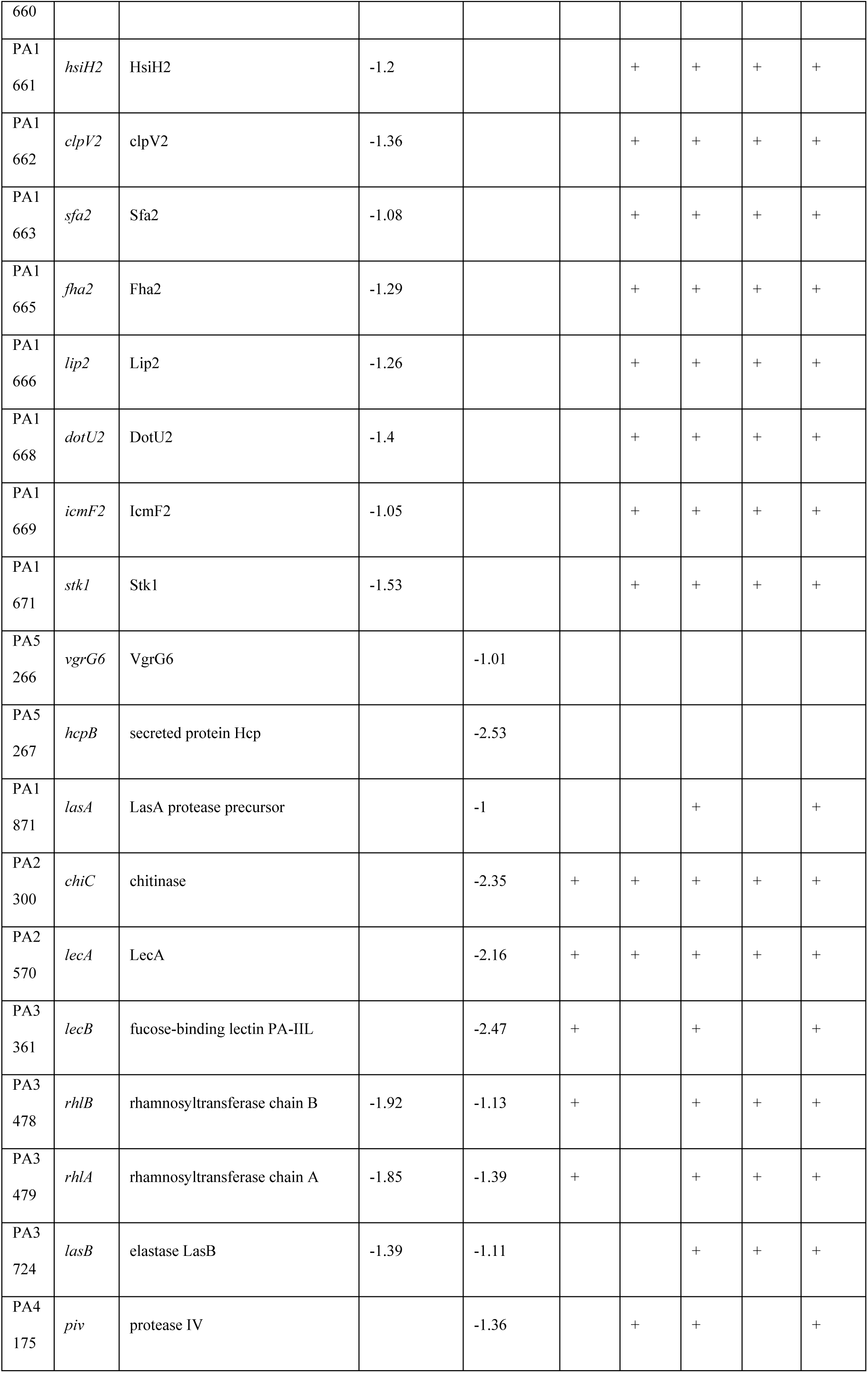
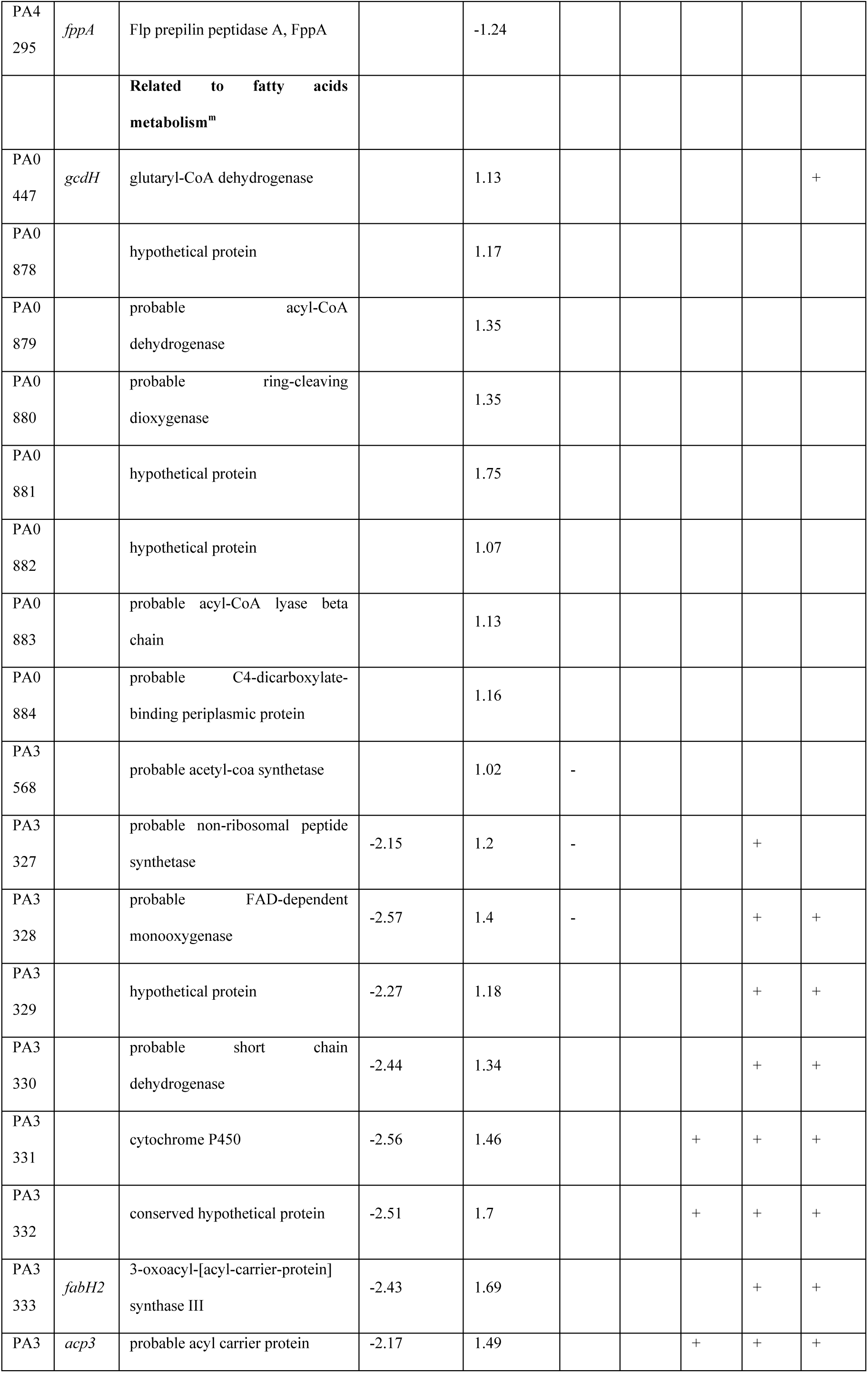
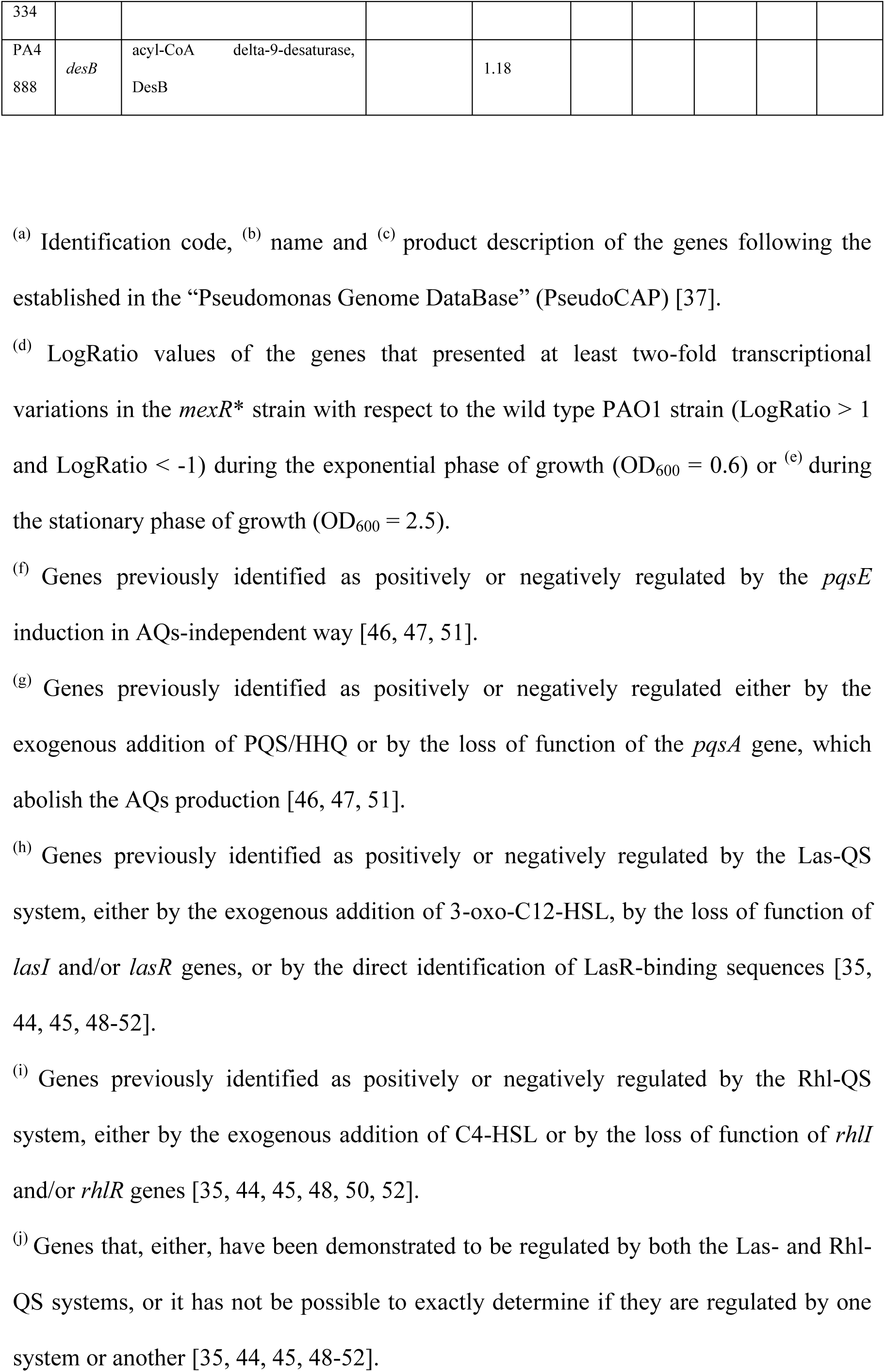
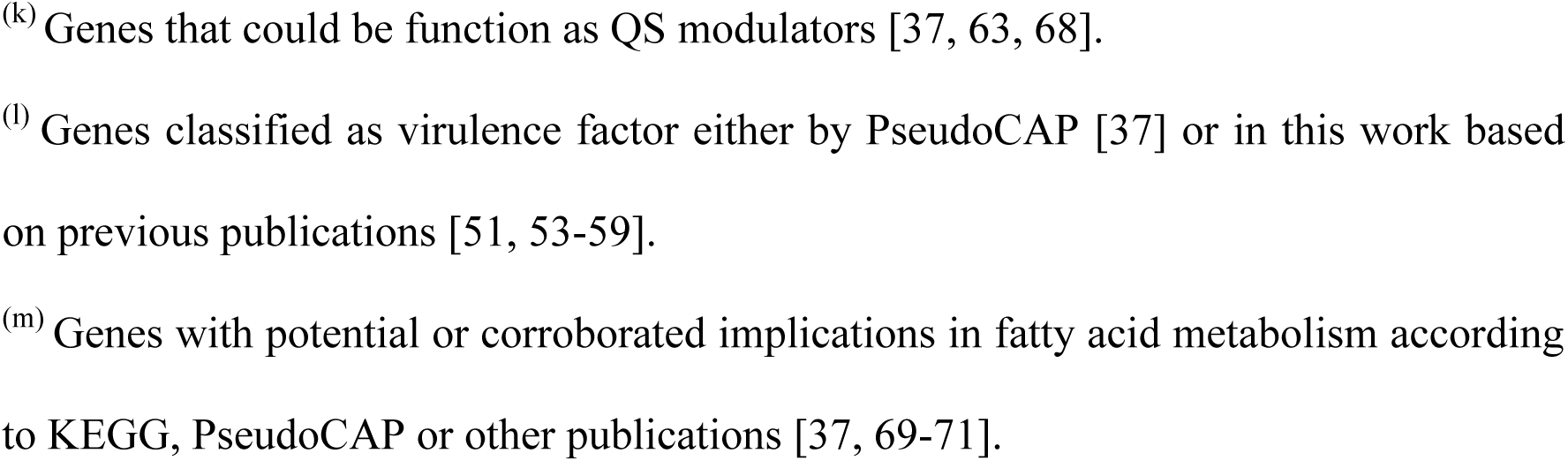
Selected list of the genes whose expression is affected by MexAB-OprM overexpression.

In order to determine which QS regulated genes present an altered expression upon MexAB overexpression, we performed a more in depth analysis of the QS regulon (Table S2), by focusing on those genes whose expression has been shown to be controlled by either PqsE, AQs or the Las and Rhl systems [35, 44–52]. As shown in Table S1, 82 of the 182 genes affected during the exponential phase of growth, and 209 out of 346 genes with transcriptional variations during the early stationary phase of growth in *mexR*\* have been associated to QS. These results evidence that overexpression of MexAB-OprM has a strong impact on QS signaling. Taking into account that the production of a large set of virulence factors is QS-regulated, it is not surprising that 76 of the 484 genes, whose expression significantly changed in the *mexR\** mutant, are catalogued as virulence factors by PseudoCAP [37] or in previously published works [51, 53–59] (Table 3 and S1). The expression of most of them, including genes of the HII-T6SS (Hcp secretion island-2 encoded type VI secretion system) and those involved in the synthesis of phenazines, pyoverdine, pyochelin, proteases, hydrogen cyanide, rhamnolipids and even QS autoinducers, was lower in the *mexR\** mutant than in the PAO1 strain (Table 3 and S1). However, the opposite was observed for other genes like some belonging to the HI-T6SS, which were expressed at higher level in the stationary growth phase in *mexR\** than in PAO1. As shown below, these results are in line with the impaired production of AQs and C4-HSL signals by the *mexR*\* mutant and the previously reported Pqs- and Rhl-mediated regulation of these virulence factors (Table S1).

To confirm the transcriptomic results, the expression of a selected set of genes was measured by RT-PCR. Since the main effect of *mexAB-OprM* overexpression was over genes belonging to the QS regulatory network, we focused our analysis in such genes. In exponential growth phase, the genes analyzed were *pqsA*, *pqsE* and *pqsH* (implicated in AQs synthesis); *rhlI* and *lasI* (responsible for C4-HSL and 3-oxo-C12-HSL synthesis, respectively); *pvdQ* (the 3-oxo-C12-HSL acylase). In the case of early stationary phase, the QS-regulated genes analyzed were *antA* (encoding the anthranilate dioxygenase large subunit), *lasB* (coding for elastase), *rhlA* (coding for the rhamnosyl transferase chain A), *lecA* (encoding the PA-I galactophilic lectin), *mexG* (encoding the subunit MexG of the RND efflux system, MexGHI-OpmD) and *phzB1* (coding for a phenazine biosynthesis protein). The RT-PCR analysis showed a similar impairment in *mexR\** levels of expression of those genes than those obtained by RNAseq (Figure 2), confirming the results obtained in the transcriptomic assays.

**Figure 2.**
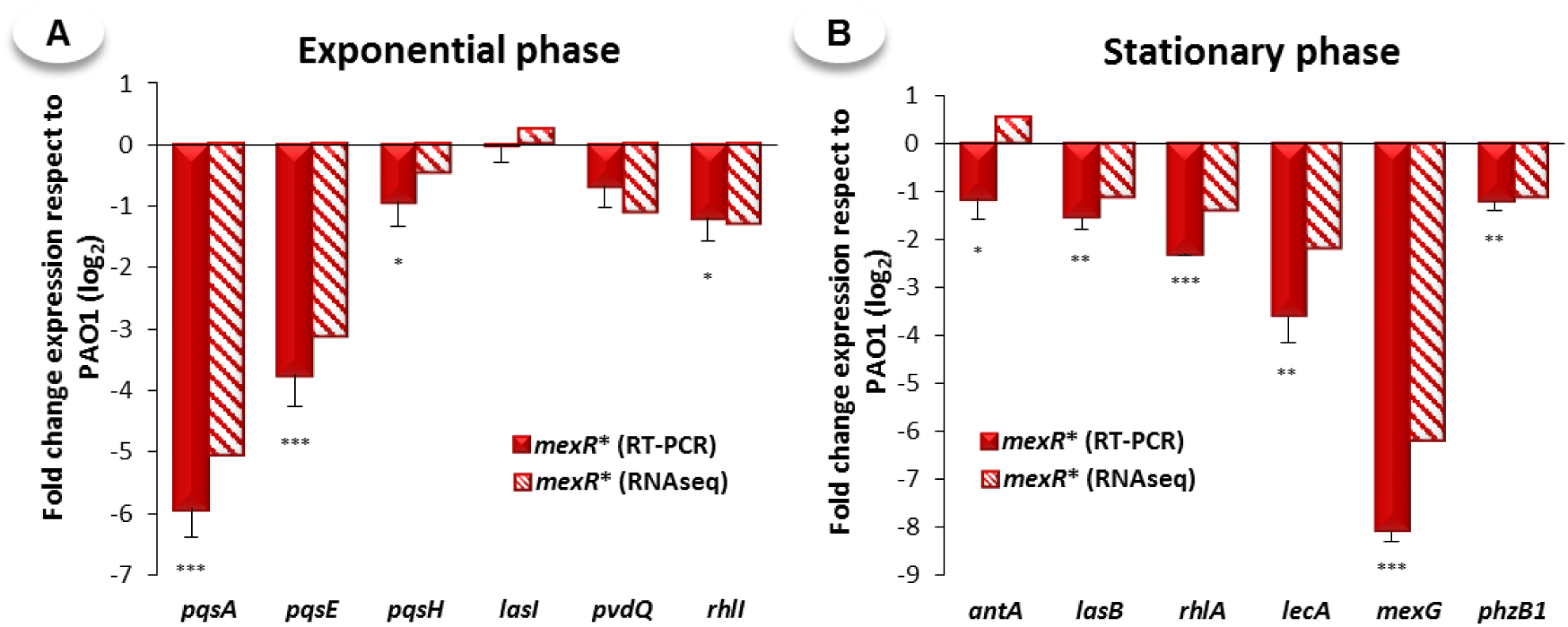
Validation of the results obtained from the transcriptomic assay in PAO1 and *mexR*\* cultures. Total RNA were extracted in triplicate for each strain at both (A) exponential (OD_600_ = 0.6) and (B) early stationary phase of growth (OD_600_ = 2.5). The expression of the genes selected for each growth phase was determined by real-time RT-PCR and compared with results obtained by RNAseq, validating the transcriptomic assays.

### The MexAB-OprM efflux system does not extrude 3-oxo-C12-HSL

It has been previously proposed that MexAB-OprM is able to extrude 3-oxo-C12-HSL [26]. We thus analyzed by LC-MS/MS the extracellular accumulation of 3-oxo-C12-HSL, C4-HSL, HHQ and PQS in PAO1 and *mexR*\* cultures grown to late stationary phase. Opposite to previous findings, 3-oxo-C12-HSL accumulation was similar in PAO1 and *mexR\**, while levels of C4-HSL and, to a greatert extent, of 2-alkyl-4-quinolones were lower when MexAB-OprM was overexpressed (Figure 3A).

**Figure 3.**
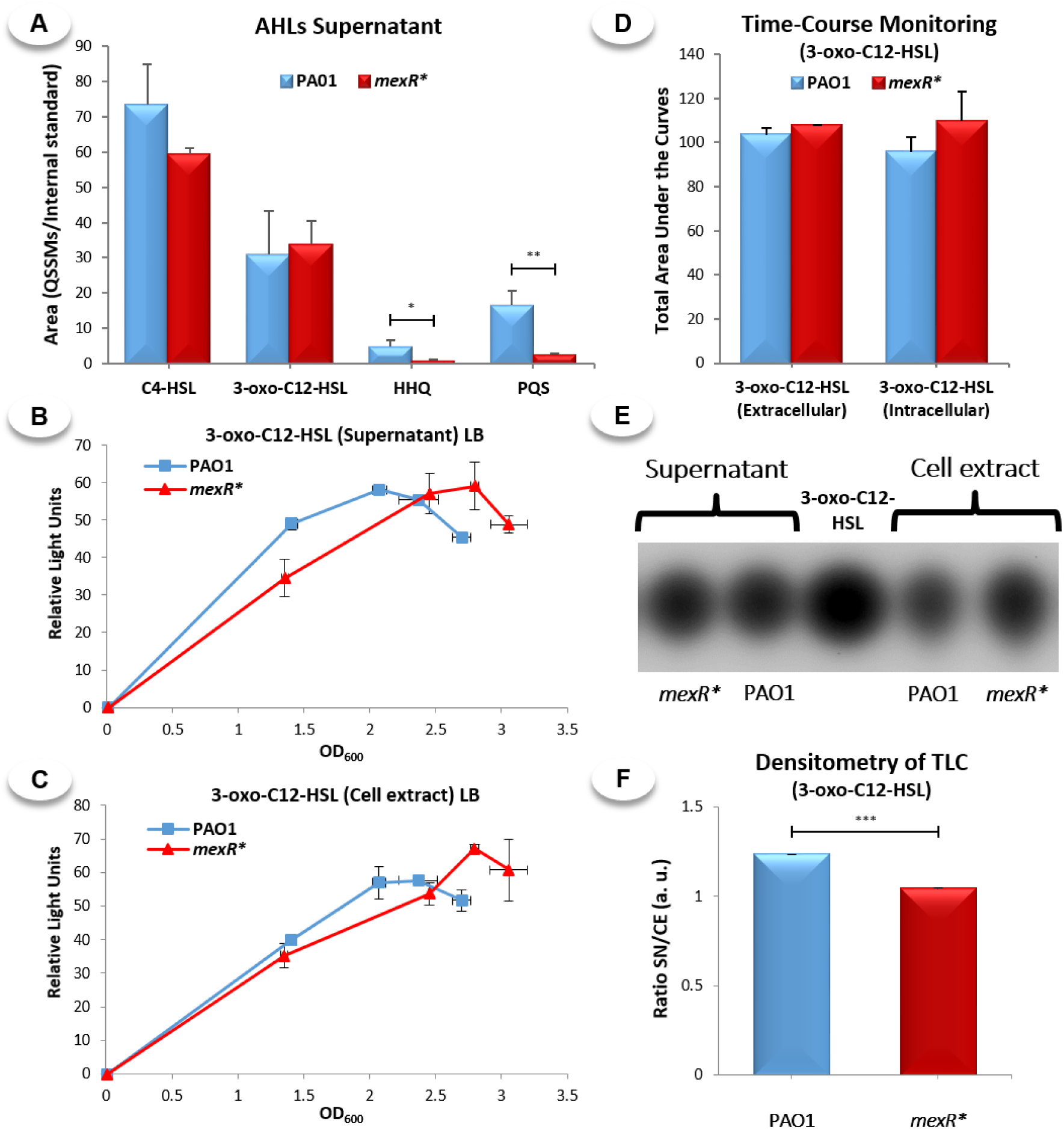
Determination of QSSMs production by PAO1 and *mexR*\* strains growing in LB medium. (A) The extracellular accumulation of C4-HSL, 3-oxo-C12-HSL, PQS and HHQ from PAO1 and *mexR*\* cultures were analyzed at late stationary phase of growth (16 hours post inoculation) by LC-MS/MS. (B) Time-course Monitoring (TCM) of 3-oxo-C12-HSL accumulation both in the supernatant and (C) inside the cells of PAO1 and *mexR*\* cultures after 4, 5, 6 and 7 hours of inoculation. (D) The area under each one of the TCM curves was calculated using “GraphPad Prism” software in order to compare the 3-oxo-C12-HSL accumulated in both PAO1 and *mexR*\* cultures along the entire incubation time. (E) Thin-Layer Chromatography (TLC) of both supernatant and cellular extracts of PAO1 and *mexR*\* cultures grown to late exponential phase (OD_600_ = 1.7) coupled to the growth of LasR-based Biosensor strain. (F) Determination of the Ratio Supernatant/Cell Extract (SN/CE) of 3-oxo-C12-HSL through densitometry analysis of the light spots derived from TLC assay. Altogether, these results demonstrate that the impaired QS response observed in the MexAB-OprM overproducer mutant, *mexR*\*, is not caused by a non-physiological extrusion of 3-oxo-C12-HSL through this efflux system.

To further analyze if MexAB-OprM is able to extrude 3-oxo-C12-HSL, the amount of this signal inside and outside PAO1 and *mexR*\* cells was determined at different growth stages. At late exponential growth phase (OD_600_ ≈ 1.2-1.8), the accumulation of 3-oxo-C12-HSL both inside and outside *mexR*\* cells was slightly lower than that of PAO1 (Figure 3B and 3C). However, once both strains reached early stationary growth phase (OD_600_ ≈ 2.5), the extracellular and intracellular accumulation of 3-oxo-C12-HSL was even slightly higher in *mexR*\* than in PAO1. To globally analyze the effect of MexAB-OprM overproduction on 3-oxo-C12-HSL accumulation, we calculated the area under each accumulation curve (Figure 3D). No significant differences were found between PAO1 and *mexR\** in any case. The ratio between the extracellular and intracellular amount of 3-oxo-C12-HSL from each strain, was also determined at late exponential phase of growth. The supernatant/cell extract (SN/CE) ratio of 3-oxo-C12-HSL in *mexR\** was even lower than to that of PAO1 (Figure 3E and 3F), which goes against an increased extrusion of this signal by the MexAB-OprM overexpressing mutant. Altogether, our results support that, opposite to what has been previously described (1999) [26], the MexAB-OprM efflux pump does not seem to extrude 3-oxo-C12-HSL, at least under our experimental conditions. An imbalance in the production of PQS, HHQ and likely C4-HSL could be the main cause for the impaired QS response associated with the MexAB-OprM overexpression.

### The production of C4-HSL is impaired in the MexAB-OprM overproducer mutant

The C4-HSL extracellular accumulation in PAO1 and *mexR*\* cultures was measured by TLC and TCM as described in Methods. Since C4-HSL freely diffuses through the plasma membrane and hence should reach an equilibrium between the extracellular and intracellular levels [26], only supernatants were measured. We found that the amount of C4-HSL was lower in *mexR\** than in PAO1 supernatantes (Figure 4A). To further support the results obtained from these measurements, the activity of the P*rhlIR* promoter, which is induced by C4-HSL in a concentration-dependent way, was determined. To do this, we generated the bioreporter strains PAO1_P*rhlI* and *mexR*\*_P*rhlI* (Table 1), which harbor the miniCTX::P*_rhlI_*-*lux* transcriptional fusion in their chromosomes. Using this combination of tools, we found that C4-HSL accumulation both outside (Figure 4A and 4B) and inside (Figure 4C) the cells was significantly lower in the *mexR*\* mutant than in the PAO1 wild type strain. Therefore, MexAB-OprM overexpression leads to a significant reduction in C4-HSL production, which could be responsible for the altered expression of some of the Rhl-regulated genes in the *mexR*\* mutant (Table 3 and S1).

**Figure 4.**
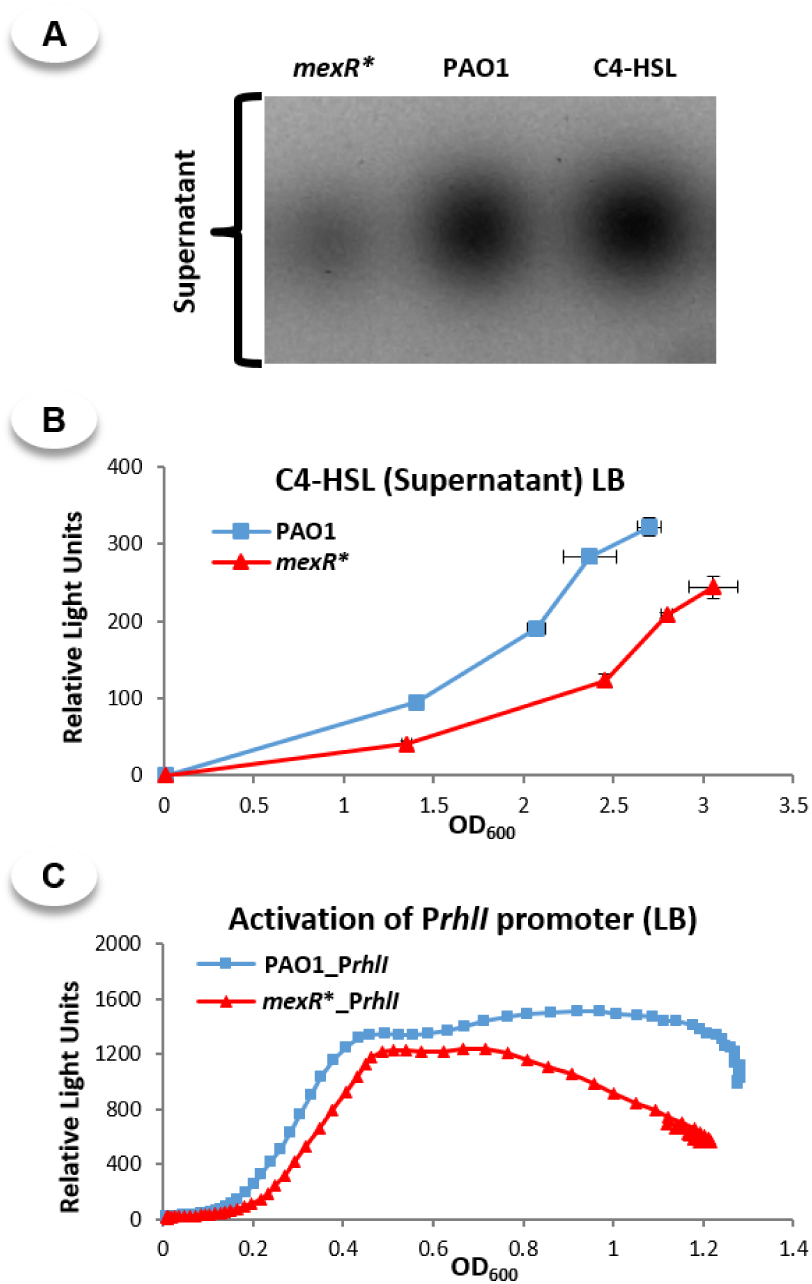
Analysis of the accumulation of C4-HSL both outside and inside the cell of PAO1 and *mexR*\* cultures grown in LB medium. (A) TLC assay of the supernatant extracts from PAO1 and *mexR*\* cultures grown to late exponential phase of growth (OD_600_ = 1.7), in which the presence of C4-HSL was revealed using the AhyR-based biosensor strain [75, 76]. (B) Analysis of the accumulation kinetics of C4-HSL in the supernatant extracts of PAO1 and *mexR*\* cultures at 4, 5, 6 and 7 hours post-inoculation. (C) Analysis of the C4-HSL-dependent activation of the P*rhlI* promoter in PAO1-P*rhlI* and *mexR*\*-P*rhlI* reporter strains along 20 hours of growth. The results showed that the production of C4-HSL is impaired in the MexAB-OprM overproducing mutant.

### The Pqs system is strongly impaired in the *mexR\** mutant

Since, both the transcriptomic analysis (Tables 3 and S1) and the LC-MS/MS results (Figure 3A) suggest that the overexpression of MexAB-OprM mainly affects Pqs-dependent regulation, we analyzed the kinetics of AQs accumulation both inside and outside bacterial cells using TLC and TCM assays coupled with the analysis of PqsR-based biosensor strains. In agreement with LC-MS/MS analysis, TLC analysis showed that overexpression of MexAB-OprM in *mexR*\* leads to a lower extracellular and intracellular accumulation of PQS and HHQ (Figure 5A and 5B). Moreover, TCM determinations indicated that this lower accumulation occurs throughout growth (Figure 5C and 5D). Altogether, our results indicate that MexAB-OprM overexpression has a strong impact on AQs production in *P. aeruginosa*, affecting their accumulation both inside and outside the cells, and, consequently, leads to a disruption of the Pqs-dependent regulation of the QS response.

**Figure 5.**
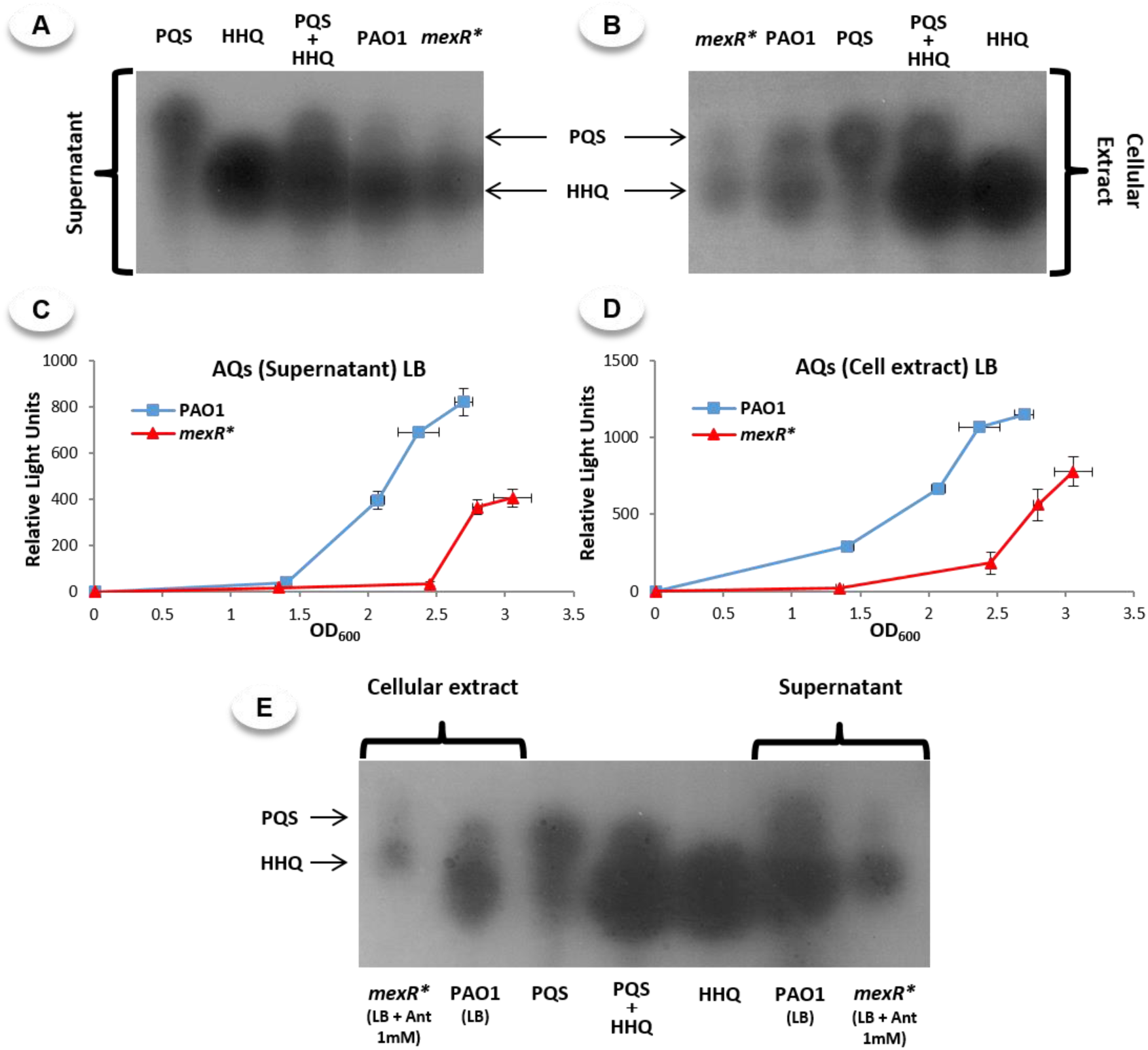
Analysis of the extracellular and cell-associated accumulation of PQS and HHQ into PAO1 and *mexR*\* cultures when grow in LB medium with or without anthranilate 1 mM. TLC assays from both (A) supernatant and (B) cellular extracts of PAO1 and *mexR*\* cultures grown to early stationary phase (OD_600_ = 2.5) in LB medium, in which the presence of PQS and HHQ were revealed using the PqsR-based biosensor strain [33, 78]. Analysis of the accumulation kinetics of AQs both (C) in the supernatant and (D) in cellular extracts of PAO1 and *mexR*\* cultures at 4, 5, 6 and 7 hours post-inoculation. (E) TLC assays from both supernatant and cellular extracts of PAO1 and *mexR*\* cultures grown to early stationary phase in LB medium supplemented with anthranilate 1 mM. These results showed that the production of PQS and HHQ is impaired in MexAB-OprM overproducing mutants and confirm that availability of anthranilate and/or its precursors is not the bottleneck for PQS and HHQ production in *mexR*\* mutant.

### The availability of anthranilate is not the bottleneck for the impaired AQs production in *mexR\**

Once demonstrated that AQs are the lowest produced QSSMs in *mexR*\*, we wanted to determine if a non-physiological MexAB-OprM-dependent extrusion of any of their precursors could be acting as a limiting step, a situation previously described for other RND overexpressing mutants in *P. aeruginosa* [27, 28]. For that purpose, *mexR\** and PAO1 cultures were supplemented with anthranilate, an immediate precursor of AQs [60, 61], and the production of PQS and HHQ was measured at early stationary growth phase. Differing to the situation reported for MexEF-OprN overproducing mutants [28], anthranilate supplementation did not restore the production of PQS and HHQ in *mexR*\* (Figure 5E). Therefore, a reduced availability of anthranilate or its precursors within the PQS/anthranilate biosynthetic pathway is not the cause for the lower PQS and HHQ production by *mexR*\*.

### Supplementation with octanoate partially restores *mexR\** QS response

Since PQS and HHQ production was not restored with anthranilate in *mexR*\*, we analyzed if the availability of octanoate, the other immediate precursor of these QSSMs, could be the limiting step in their production. Since it has been described that the production of both AQs and pyocyanin increase in presence of this compound [31], the kinetics of extracellular and intracellular AQs accumulation throughout the cell-growth of both PAO1 and *mexR*\* strains grown in LB supplemented with octanoate 5mM was determined by TCM. In these conditions, production of AQs was delayed in *mexR*\* when compared with PAO1 (Figure 6A and 6B). However, once the stationary phase of growth is reached (OD_600_ > 2.5), AQs production in *mexR*\* and PAO1 is similar, being the accumulation in the supernatant even slightly higher in the MexAB-OprM overexpressing mutant. Furthermore, the TLC analysis of extracellular and intracellular accumulation of PQS and HHQ at 7 hours post-inoculation (Figure 6C) further supports that supplementation with octanoate restored almost fully PQS/HHQ production in *mexR\**.

**Figure 6.**
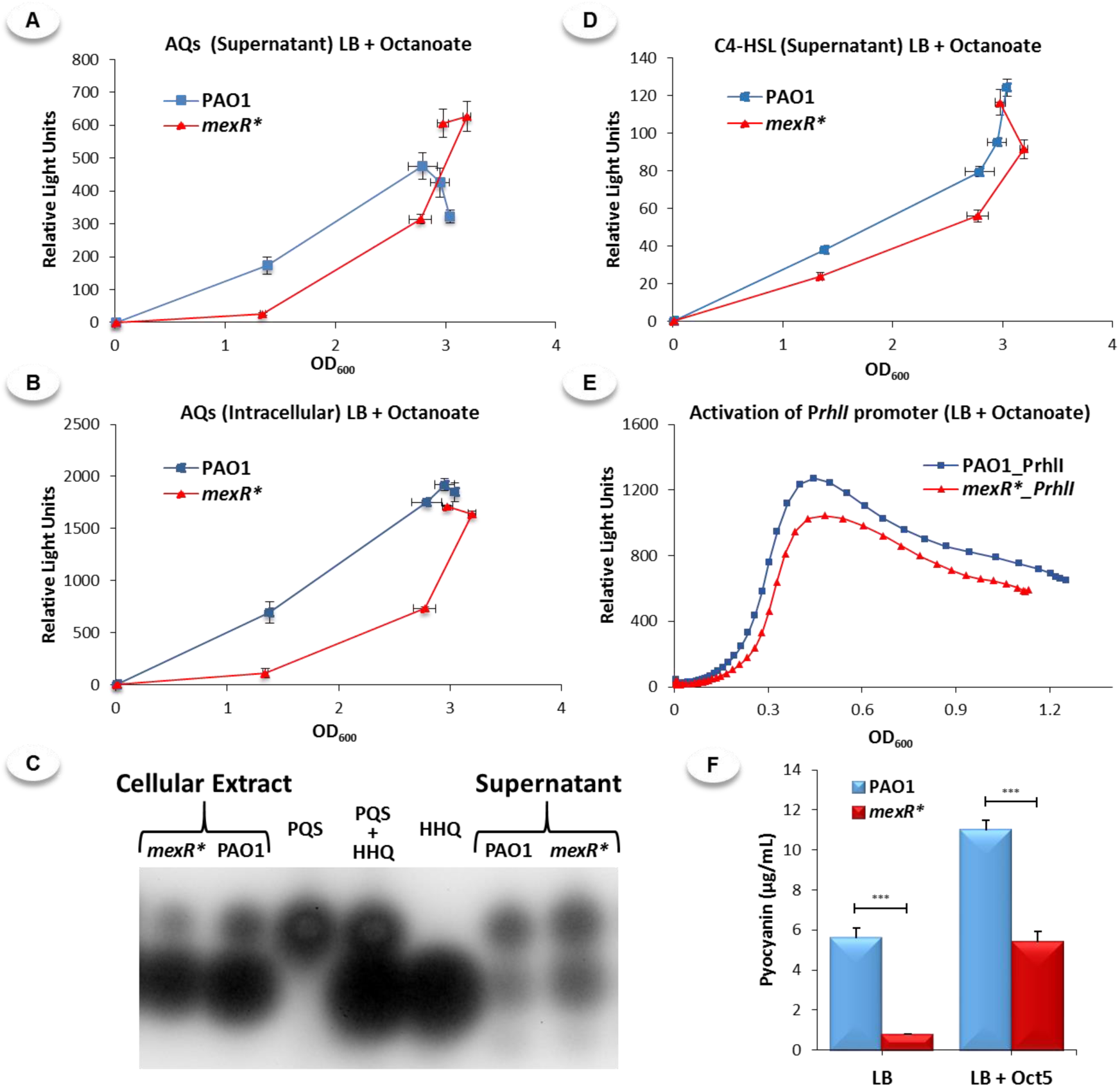
Analysis of the production of PQS, HHQ, C4-HSL and pyocyanin by PAO1 and *mexR*\* cultures grown in LB medium supplemented with octanoate 5 mM. Analysis by TCM of (A and B) AQs and (D) C4-HSL accumulation both (A and D) outside and (B) inside the cells of PAO1 and *mexR*\* cultures after 4, 5, 6 and 7 hours of incubation. (C) TLC of supernatant and cellular extracts from cultures incubated along 7 hours in LB medium supplemented with octanoate 5 mM, which was revealed using PqsR-based biosensor strain [33, 78]. (E) Real-time monitoring of the C4-HSL-dependent activation of the P*rhlI* promoter in PAO1 and *mexR*\* cultures grown along 20 hours in presence of octanoate 5 mM. (F) Quantification of pyocyanin production by PAO1 and *mexR*\* strains grown along 20 hours in LB medium supplemented with octanoate 5 mM. The results showed that LB supplementation with octanoate 5 mM partially restore the production of AQs, C4-HSL and pyocyanin in MexAB-OprM overproducer mutant, suggesting that a decreased availability of octanoate or any of its precursors should be the cause of the QS impairment observed in *mexR*\*.

Once we determined that octanoate supplementation restores AQs production in *mexR*\*, we analyzed whether or not the presence of octanoate may also allow the recovering of C4-HSL levels, which were also significantly affected in *mexR*\* (Figure 4). As shown in Figures 6D and 6E, octanoate supplementation affects C4-HSL production by *mexR*\* in a similar way than for AQs, keeping a delay in C4-HSL accumulation at early growth time, but reaching C4-HSL levels close to those observed in PAO1 once stationary phase is reached. To get a functional validation of the effect of octanoate supplementation on the recovery of the QS response in *mexR\**, the production of pyocyanin by PAO1 and *mexR*\* growing in presence or absence of octanoate was measured. In absence of octanoate, the *mexR*\* strain produced less than 15% the amount of pyocyanin produced by the wild type strain, while this production in presence of octanoate reached to 50% of the amount produced by PAO1 (Figure 6F).

The almost total recovery of AQs and C4-HSL production, together with the partial recovery of pyocyanin production, prompted by octanoate supplementation, suggests that the QS defects associated to the MexAB-OprM overexpression should be due, at least in part, to a decrease in the intracellular availability of this immediate precursor of AQs.

## Discussion

One of the key features for the successful adaptation of bacteria to the continuous changing environment lies in the plasticity of their physiology and in the presence of global regulation networks able to translate environmental signals to the cells [10, 62]. Although bacteria present a wide variety of classical master regulators, modulation of the activity of such regulatory networks can be also achieved through the action of other elements involved in fundamental aspects of bacterial physiology [63–65]. One of the ways for achieving such modulation is by interfering with the available concentrations of the signals that trigger the regulatory networks. For this purpose, multidrug efflux pumps can be particularly well suited, because the activity of these antibiotic resistance determinants can affect the bacterial physiology through the extrusion of endogenous/exogenous molecular compounds with relevance for bacterial physiology [3, 7]. As a consequence of this extrusion, the expression of a large number of genes, including those encoding virulence determinants, may be altered, which has been usually considered as the fitness cost associated to the acquisition of antibiotic resistance. However, the fact that the expression of these efflux systems can be triggered by specific signals/environmental conditions, together with supporting evidence for the impact of their increased expression on specific bacterial processes, suggests that these collateral effects on bacterial physiology may have adaptive values.

One of the regulation processes where efflux pumps may have a role in *P. aeruginosa* is the QS response. Notably, different efflux systems seem to modulate this regulatory network, a feature that at a first sight seem to be a non-needed redundant function. The first one to be studied was MexAB-OprM and the proposed mechanism of modulating the QS response was the active extrusion the QS signal 3-oxo-C12-HSL through this efflux system [26]. Later on, MexCD-OprJ and MexEF-OprN overexpression was shown to downregulate some QS-controlled phenotypes [27, 28]. Notably, although it has been shown that both efflux systems are able to extrude kynurenine and HHQ [27, 28, 66], both biosynthetic precursors of PQS [67], there were differences in the subjacent cause of impaired QS response associated with MexCD-OprJ overexpression (mainly HHQ extrusion) or with the increased MexEF-OprN expression (mainly kynurenine efflux) [27, 28]. Concerning MexGHI-OpmD, it has been proposed that this efflux system is able to extrude anthranilate and 5-methylphenazine-1-carboxylate (5-Me-PCA), which are immediate precursors of AQs and pyocyanin, respectively, being the latter a known virulence factor regulated by QS response [29, 53].

Given this apparent functional redundancy, we wanted to get more information on the mechanism leading to the impaired expression of QS-regulated genes in a MexAB-OprM overexpressing mutant. Although it was earlier stated that MexAB-OprM extrudes 3-oxo-C12-HSL [26], our data do not support that the putative non-physiological extrusion of this AHL by MexAB-OprM is the cause of the impaired QS response of *mexR\**. Our results, however, showed that production of PQS/HHQ, and to a lesser extent, of C4-HSL were significantly lower in a MexAB-OprM overexpressing mutant that in the wild-type strain, while 3-oxo-C12-HSL production and intracellular accumulation did not change significantly. These results are in line with our transcriptomic data, which showed that the expression of the genes responsible for PQS/HHQ biosynthesis and some of the main AQs-regulated genes were the most affected in the *mexR*\* mutant. Altogether, we concluded that, opposite to previously published work [26], the impaired QS response of the MexAB-OprM overproducing mutant is mainly caused by the low production of PQS and HHQ, rather than by a non-physiological extrusion of 3-oxo-C12-HSL through this multidrug efflux pump.

The observed low production of AQs in the MexAB-OprM overexpressing mutant could be due to a reduced availability of their metabolic precursors as it has been previously described in the case of mutants overexpressing either MexEF-OprN or MexCD-OprJ [27, 28]. We have seen that supplementation of *mexR*\* cultures with anthranilate, one of the immediate precursors of HHQ and PQS [60, 61], did not restore the wild-type levels of these QSSMs (Figure 5E). However, supplementation with octanoate, the other immediate precursor of PQS and HHQ [31], allowed almost full restoration of AQs production in *mexR*\* at early stationary growth phase. Further, octanoate supplementation had similar effects on C4-HSL production by *mexR*\*, resulting in an increase of this AHL in the MexAB-OprM overexpressing mutant to similar levels to those observed in the PAO1 wild type strain. Altogether our results suggest that the main cause for the impaired production of PQS/HHQ observed in MexAB-OprM overproducing mutants is the low intracellular availability of octanoate. In addition, we propose that this defect in AQs synthesis is also responsible for the low production of C4-HSL in *mexR*\* by decreasing the Pqs-dependent expression of *rhlI* and *acp1* genes, which are implicated in the synthesis of this AHL (Table 3) [68]. In agreement with a role of MexAB-OprM in octanoate trafficking/metabolism, we found that MexAB-OprM overexpression led to changes in the expression of many genes involved in fatty acids metabolism such as *fabH2*, *desB* or *gcdH*, as well as some lipoproteins like *acp1* or *acp3* [37, 69–71]. Taking this information into consideration, we propose that an altered fatty acids metabolism in the MexAB-OprM overproducer mutant, *mexR*\*, which may lead to changes on intracellular availability of octanoic acid, the immediate precursor of PQS and HHQ, could be a main reason for the strong decreased production of these QSSMs observed in this mutant. Furthermore, taking into account that, even in the presence of octanoate, the *mexR*\* mutant presented a delay in AQs and C4-HSL biosynthesis with respect to PAO1, we cannot discard that some additional factors may also be affecting the production of QSSMs and the activation of the QS response. A slight extrusion of PQS and/or HHQ through MexAB-OprM might be involved in the observed phenotype as well, since the overproducer mutant, in presence of octanoate, accumulates more AQs outside the cells with respect to the parental PAO1 (Figure 6A and 6C), whereas the opposite trend was observed in cellular extracts (Figure 6B and 6C).

This work, together with other previous studies [27-29, 53, 66], show that, although the consequences of overexpressing each multidrug efflux pump are similar (an impaired QS response), the underlying mechanism by which the overexpression of these RND efflux systems interferes with the *P. aeruginosa* QS response is different in every case (Figure 7). Furthermore, they support the concept that *P. aeruginosa* might modulate the QS response by regulating the expression of several RND efflux systems in response to nutritional and environmental signals/cues (Figure 7).

**Figure 7.**
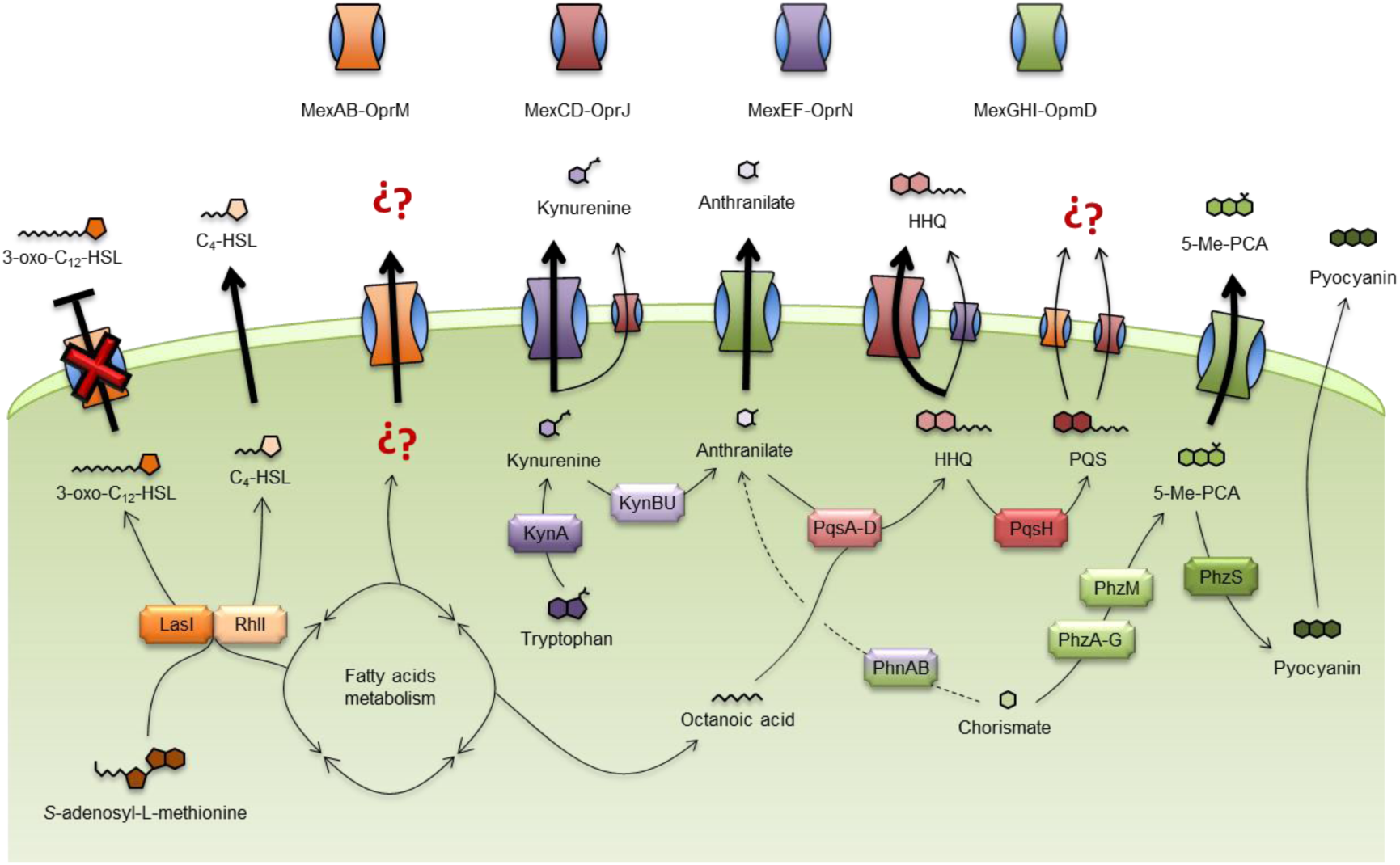
RND-dependent production of the QS signals in *P. aeruginosa*. The main autoinducer signals involve in the QS response in *P. aeruginosa* are 3-oxo-C12-HSL, C4-HSL, HHQ and PQS [10]. The synthesis of the two homoserine lactones, 3-oxo-C12-HSL and C4-HSL, are carried out by the synthases enzyme LasI and RhlI, respectively, which use both acyl carrier proteins and *S*-adenosyl-L-methionine as precursors [68, 80]. The synthesis of HHQ is mediated by the synthetic enzymes PqsA, PqsB, PqsC and PqsD using anthranilate and octanoic acid as immediate precursors, while PQS is a chemical modification of HHQ carried out by PqsH [31]. Pearson *et al.* (1999) [26] described that C4-HSL is an autoinducer signal able to freely diffuse through the plasma membrane and proposed that 3-oxo-C12-HSL could be actively extruded through MexAB-OprM efflux system. However, in this work we demonstrate that MexAB-OprM is not able to extrude this QS signal. At the same time, we suggest that the overexpression of MexAB-OprM produces an imbalance in the fatty acid metabolism, which in last instance could be causing the low intracellular availability of octanoate by which this antibiotic resistant mutant present defects in PQS/HHQ production and an impaired Pqs-dependent regulation of the QS response. In addition, the MexCD-OprJ and MexEF-OprN efflux systems are able to extrude both kynurenine and HHQ [27, 28, 66], which are both precursors of PQS [67]. However, the affinity by one or another substrate seems to be different, since the bottleneck for the impaired PQS production observed in the MexCD-OprJ overproducer mutants is the massive extrusion of HHQ [27], while in MexEF-OprN overproducer mutants is the massive extrusion of kynurenine [28]. Furthermore, it has been suggested that MexAB-OprM and MexCD-OprJ could be also extruding, to a lesser extent, PQS, since it has been observed a slight increase on outside/cell-associated PQS accumulating ratio respect to PAO1 wild type strain (This work and [27]). Finally, it has been described that MexGHI-OpmD is able to extrude anthranilate and 5-methylphenazine-1-carboxylate (5-Me-PCA), which are either precursors of PQS and of the QS-regulated virulence factor, pyocyanin, respectively [29, 53].

Whereas redundancy may help in keeping the robustness of living systems, it also bears a cost and it is sometimes difficult to foresee the reason of this redundancy. Concerning the potential modulation of the QS response through the activity of different efflux pumps, the fact that the molecular basis of such modulation are different in each case support that the role of these QS-related efflux systems should not be considered strictly redundant. For that reason, we propose the term ‘apparent redundancy’ to define this kind of situation in which similar phenotypes, produced by genes belonging to the same family (in our case RND efflux pumps) are prompted by different mechanisms. This situation may allow to keep both the homeostasis and the plasticity of physiological networks with a fundamental role in bacterial adaptation to habitat changes as the QS response. Indeed, the fact that the QS network of *P. aeruginosa* presents several interconnected control levels that, as here reported, can also be modulated at different checkpoints *via* changing the level of MDR efflux pumps expression, indicates that this is a robust network (redundant) but also able of responding to different signals/cues (’apparently redundant’). One example of that RND-mediated re-accommodation of the QS response is the work published by Oshri *et al.* (2018) [72], in which the authors shown that a *lasR*-null mutant, with a defective QS response, is able to regain a partial QS-dependent cooperative behavior when grew in casein medium (QS-favored conditions). The study revealed that this partial regained of casein growth was associated with an increased activation of the RhlIR system mediated by the reduced activity of MexEF-OprN, which was in turn caused by selection of non-functional mutation of *mexT* that encodes a master regulator needed for *mexEF-oprN* expression [73]. These evidences further reinforce the role that efflux pumps have in the non-canonical modulation of the *P. aeruginosa* QS response.

Our study and those previously published have shown that the expression of RND efflux systems can affect the QS regulatory network at different levels (Figure 7). However, the original role of these multidrug efflux pumps on the modulation of the QS response, considering the environmental and nutritional bacterial requirements remains unknown, particularly if we take into consideration that the physiological signals that trigger the expression of these efflux systems are largely ignored. For example, besides their role in modulating the QS response in clonal populations, the activity of these RND systems could have unknown social implications in heterogeneous populations of *P. aeruginosa*. In environments where the acquisition of required public goods is energetically expensive, the emergence of cheaters unable to respond to QSSMs but able to benefit from cooperative behaviors, is common [12, 19]. In this context, the expression level of each of the QS-related RND systems could determine the ability of *P. aeruginosa* to act, in function of the needs, either as social cooperators or as social cheaters.

Altogether, this work, along with others previously published, evidences the complexity of the relationship between several RND efflux systems and the QS regulatory network. Further, we propose that the “apparent redundancy” observed among different efflux systems belonged to a particular bacterial species could have an adaptive role on its successful colonization of the range of niches where it can be present.

## Acknowledgments

Work in the laboratory of JLM has been supported by grants from the Instituto de Salud Carlos III (Spanish Network for Research on Infectious Diseases [RD16/0016/0011]), from the Spanish Ministry of Economy, Industry and Competitivity (BIO2017-83128-R) and from the Autonomous Community of Madrid (B2017/BMD-3691). This work was also supported by funding from the Biotechnology and Biological Sciences Research Council (Award Number BB/R012415/1). MC is partly funded by the National Biofilms Innovation Centre (NBIC) with is an Innovation and Knowledge Centre funded by the Biotechnology and Biological Sciences Research Council, InnovateUK and Hartree Centre. Thanks are given to Stephan Heeb for providing the *E. coli* S17 with the miniCTX::P*_rhlI_*-*lux* construction. The funders had no role in study design, data collection and interpretation, or the decision to submit the work for publication.

## Supplementary material

**Table S1.** Complete list of genes whose expression presented at least two-fold change in the *mexR*\* strain with respect to the wild type PAO1 strain (LogRatio > 1 and LogRatio < −1) during the exponential (OD_600_ = 0.6) or stationary phase of growth (OD_600_ = 2.5)

**Table S2.** Review of the genes whose expression is regulated in a QS-dependent way, making difference between each one of the QS system implicated in the control of their expression level.

## Bibliography

1. WHO, Global Priority List of Antibiotic-Resistant Bacteria to Guide Research, Discovery, and Development of New Antibiotics. World Health Organization: Essential medicines and health products, 2017: p. 7.

2. Driscoll, J.A., S.L. Brody, and M.H. Kollef, The epidemiology, pathogenesis and treatment of Pseudomonas aeruginosa infections. Drugs, 2007. 67(3): p. 351–68.

3. Blanco, P., et al., Bacterial Multidrug Efflux Pumps: Much More Than Antibiotic Resistance Determinants. Microorganisms, 2016. 4(1).

4. Olivares, J., et al., The intrinsic resistome of bacterial pathogens. Front Microbiol, 2013. 4: p. 103.

5. Blair, J.M., G.E. Richmond, and L.J. Piddock, Multidrug efflux pumps in Gram-negative bacteria and their role in antibiotic resistance. Future Microbiol, 2014. 9(10): p. 1165–77.

6. Hernando-Amado, S., et al., Multidrug efflux pumps as main players in intrinsic and acquired resistance to antimicrobials. Drug Resist Updat, 2016. 28: p. 13–27.

7. Alcalde-Rico, M., et al., Multidrug Efflux Pumps at the Crossroad between Antibiotic Resistance and Bacterial Virulence. Front Microbiol, 2016. 7: p. 1483.

8. Morita, Y., et al., Roles of MexXY- and MexAB-multidrug efflux pumps in intrinsic multidrug resistance of Pseudomonas aeruginosa PAO1. J Gen Appl Microbiol, 2001. 47(1): p. 27–32.

9. Chalhoub, H., et al., Loss of activity of ceftazidime-avibactam due to Mex-AB-OprM efflux and overproduction of AmpC cephalosporinase in Pseudomonas aeruginosa isolated from patients suffering from cystic fibrosis. Int J Antimicrob Agents, 2018.

10. Williams, P. and M. Camara, Quorum sensing and environmental adaptation in Pseudomonas aeruginosa: a tale of regulatory networks and multifunctional signal molecules. Curr Opin Microbiol, 2009. 12(2): p. 182–91.

11. Miller, M.B. and B.L. Bassler, Quorum sensing in bacteria. Annu Rev Microbiol, 2001. 55: p. 165–99.

12. Wilder, C.N., S.P. Diggle, and M. Schuster, Cooperation and cheating in Pseudomonas aeruginosa: the roles of the las, rhl and pqs quorum-sensing systems. ISME J, 2011. 5(8): p. 1332–43.

13. Vidal, J.E., et al., The LuxS-dependent quorum-sensing system regulates early biofilm formation by Streptococcus pneumoniae strain D39. Infect Immun, 2011. 79(10): p. 4050–60.

14. Holm, A., K.E. Magnusson, and E. Vikstrom, Pseudomonas aeruginosa N-3-oxo-dodecanoyl-homoserine Lactone Elicits Changes in Cell Volume, Morphology, and AQP9 Characteristics in Macrophages. Front Cell Infect Microbiol, 2016. 6: p. 32.

15. Pearson, J.P., E.C. Pesci, and B.H. Iglewski, Roles of Pseudomonas aeruginosa las and rhl quorum-sensing systems in control of elastase and rhamnolipid biosynthesis genes. J Bacteriol, 1997. 179(18): p. 5756–67.

16. Heurlier, K., V. Denervaud, and D. Haas, Impact of quorum sensing on fitness of Pseudomonas aeruginosa. Int J Med Microbiol, 2006. 296(2-3): p. 93–102.

17. Gupta, R. and M. Schuster, Negative regulation of bacterial quorum sensing tunes public goods cooperation. ISME J, 2013. 7(11): p. 2159–68.

18. Kostylev, M., et al., Evolution of the Pseudomonas aeruginosa quorum-sensing hierarchy. Proc Natl Acad Sci U S A, 2019. 116(14): p. 7027–7032.

19. Feltner, J.B., et al., LasR Variant Cystic Fibrosis Isolates Reveal an Adaptable Quorum-Sensing Hierarchy in Pseudomonas aeruginosa. MBio, 2016. 7(5).

20. Choi, Y., et al., Growth phase-differential quorum sensing regulation of anthranilate metabolism in Pseudomonas aeruginosa. Mol Cells, 2011. 32(1): p. 57–65.

21. Mellbye, B. and M. Schuster, Physiological framework for the regulation of quorum sensing-dependent public goods in Pseudomonas aeruginosa. J Bacteriol, 2014. 196(6): p. 1155–64.

22. Schafhauser, J., et al., The stringent response modulates 4-hydroxy-2-alkylquinoline biosynthesis and quorum-sensing hierarchy in Pseudomonas aeruginosa. J Bacteriol, 2014. 196(9): p. 1641–50.

23. Kang, H., et al., Crystal structure of Pseudomonas aeruginosa RsaL bound to promoter DNA reaffirms its role as a global regulator involved in quorum-sensing. Nucleic Acids Res, 2017. 45(2): p. 699–710.

24. Deziel, E., et al., The contribution of MvfR to Pseudomonas aeruginosa pathogenesis and quorum sensing circuitry regulation: multiple quorum sensing-regulated genes are modulated without affecting lasRI, rhlRI or the production of N-acyl-L-homoserine lactones. Mol Microbiol, 2005. 55(4): p. 998–1014.

25. Evans, K., et al., Influence of the MexAB-OprM multidrug efflux system on quorum sensing in Pseudomonas aeruginosa. J Bacteriol, 1998. 180(20): p. 5443–7.

26. Pearson, J.P., C. Van Delden, and B.H. Iglewski, Active efflux and diffusion are involved in transport of Pseudomonas aeruginosa cell-to-cell signals. J Bacteriol, 1999. 181(4): p. 1203–10.

27. Alcalde-Rico, M., et al., Role of the Multidrug Resistance Efflux Pump MexCD-OprJ in the Pseudomonas aeruginosa Quorum Sensing Response. Front Microbiol, 2018. 9: p. 2752.

28. Olivares, J., et al., Overproduction of the multidrug efflux pump MexEF-OprN does not impair Pseudomonas aeruginosa fitness in competition tests, but produces specific changes in bacterial regulatory networks. Environ Microbiol, 2012. 14(8): p. 1968–81.

29. Aendekerk, S., et al., The MexGHI-OpmD multidrug efflux pump controls growth, antibiotic susceptibility and virulence in Pseudomonas aeruginosa via 4-quinolone-dependent cell-to-cell communication. Microbiology, 2005. 151(Pt 4): p. 1113–25.

30. Sanchez, P., et al., Fitness of in vitro selected Pseudomonas aeruginosa nalB and nfxB multidrug resistant mutants. J Antimicrob Chemother, 2002. 50(5): p. 657–64.

31. Dulcey, C.E., et al., The end of an old hypothesis: the pseudomonas signaling molecules 4-hydroxy-2-alkylquinolines derive from fatty acids, not 3-ketofatty acids. Chem Biol, 2013. 20(12): p. 1481–91.

32. Yates, E.A., et al., N-Acylhomoserine Lactones Undergo Lactonolysis in a pH-, Temperature-, and Acyl Chain Length-Dependent Manner during Growth of Yersinia pseudotuberculosis and Pseudomonas aeruginosa. Infection and Immunity, 2002. 70(10): p. 5635–5646.

33. Fletcher, M.P., et al., Biosensor-based assays for PQS, HHQ and related 2-alkyl-4-quinolone quorum sensing signal molecules. Nat Protoc, 2007. 2(5): p. 1254–62.

34. Essar, D.W., et al., Identification and characterization of genes for a second anthranilate synthase in Pseudomonas aeruginosa: interchangeability of the two anthranilate synthases and evolutionary implications. J Bacteriol, 1990. 172(2): p. 884–900.

35. Schuster, M., et al., Identification, Timing, and Signal Specificity of Pseudomonas aeruginosa Quorum-Controlled Genes: a Transcriptome Analysis. Journal of Bacteriology, 2003. 185(7): p. 2066–79.

36. Sultan, M., et al., A simple strand-specific RNA-Seq library preparation protocol combining the Illumina TruSeq RNA and the dUTP methods. Biochem Biophys Res Commun, 2012. 422(4): p. 643–6.

37. Winsor, G.L., et al., Enhanced annotations and features for comparing thousands of Pseudomonas genomes in the Pseudomonas genome database. Nucleic Acids Res, 2016. 44(D1): p. D646–53.

38. Charles D. Warden, Y.-C.Y., and Xiwei Wu, Optimal Calculation of RNA-Seq Fold-Change Values. Int J Comput Bioinfo In Silico Model, 2013. 2(6): p. 285–292.

39. Livak, K.J. and T.D. Schmittgen, Analysis of relative gene expression data using real-time quantitative PCR and the 2(-Delta Delta C(T)) Method. Methods, 2001. 25(4): p. 402–8.

40. Ortori, C.A., et al., Simultaneous quantitative profiling of N-acyl-L-homoserine lactone and 2-alkyl-4(1H)-quinolone families of quorum-sensing signaling molecules using LC-MS/MS. Anal Bioanal Chem, 2011. 399(2): p. 839–50.

41. Hoang, T.T., et al., Integration-proficient plasmids for Pseudomonas aeruginosa: site-specific integration and use for engineering of reporter and expression strains. Plasmid, 2000. 43(1): p. 59–72.

42. Maseda, H., et al., Enhancement of the mexAB-oprM efflux pump expression by a quorum-sensing autoinducer and its cancellation by a regulator, MexT, of the mexEF-oprN efflux pump operon in Pseudomonas aeruginosa. Antimicrob Agents Chemother, 2004. 48(4): p. 1320–8.

43. Evans, K. and K. Poole, The MexA-MexB-OprM multidrug efflux system of Pseudomonas aeruginosa is growth-phase regulated. FEMS Microbiol Lett, 1999. 173(1): p. 35–9.

44. Wagner, V.E., R.J. Gillis, and B.H. Iglewski, Transcriptome analysis of quorum-sensing regulation and virulence factor expression in Pseudomonas aeruginosa. Vaccine, 2004. 22 **Suppl 1**: p. S15–20.

45. Wagner, V.E., et al., Microarray Analysis of Pseudomonas aeruginosa Quorum-Sensing Regulons: Effects of Growth Phase and Environment. J Bacteriol, 2003. 185(7): p. 2080–2095.

46. Rampioni, G., et al., Unravelling the Genome-Wide Contributions of Specific 2-Alkyl-4-Quinolones and PqsE to Quorum Sensing in Pseudomonas aeruginosa. PLoS Pathog, 2016. 12(11): p. e1006029.

47. Rampioni, G., et al., Transcriptomic analysis reveals a global alkyl-quinolone-independent regulatory role for PqsE in facilitating the environmental adaptation of Pseudomonas aeruginosa to plant and animal hosts. Environ Microbiol, 2010. 12(6): p. 1659–73.

48. Hentzer, M., et al., Attenuation of Pseudomonas aeruginosa virulence by quorum sensing inhibitors. EMBO J, 2003. 22(15): p. 3803–15.

49. Gilbert, K.B., et al., Global position analysis of the Pseudomonas aeruginosa quorum-sensing transcription factor LasR. Mol Microbiol, 2009. 73(6): p. 1072–85.

50. Schuster, M. and E.P. Greenberg, Early activation of quorum sensing in Pseudomonas aeruginosa reveals the architecture of a complex regulon. BMC Genomics, 2007. 8: p. 287.

51. Lesic, B., et al., Quorum sensing differentially regulates Pseudomonas aeruginosa type VI secretion locus I and homologous loci II and III, which are required for pathogenesis. Microbiology, 2009. 155(Pt 9): p. 2845–55.

52. Sana, T.G., et al., The second type VI secretion system of Pseudomonas aeruginosa strain PAO1 is regulated by quorum sensing and Fur and modulates internalization in epithelial cells. J Biol Chem, 2012. 287(32): p. 27095–105.

53. Sakhtah, H., et al., The Pseudomonas aeruginosa efflux pump MexGHI-OpmD transports a natural phenazine that controls gene expression and biofilm development. Proc Natl Acad Sci U S A, 2016. 113(25): p. E3538–47.

54. Russell, A.B., et al., Type VI secretion delivers bacteriolytic effectors to target cells. Nature, 2011. 475(7356): p. 343–7.

55. Chen, L., et al., Composition, function, and regulation of T6SS in Pseudomonas aeruginosa. Microbiol Res, 2015. 172: p. 19–25.

56. Ball, G., et al., A novel type II secretion system in Pseudomonas aeruginosa. Mol Microbiol, 2002. 43(2): p. 475–85.

57. Pelzer, A., et al., Subtilase SprP exerts pleiotropic effects in Pseudomonas aeruginosa. Microbiologyopen, 2014. 3(1): p. 89–103.

58. Lee, X., et al., Identification of the biosynthetic gene cluster for the Pseudomonas aeruginosa antimetabolite L-2-amino-4-methoxy-trans-3-butenoic acid. J Bacteriol, 2010. 192(16): p. 4251–5.

59. Folders, J., et al., Characterization of Pseudomonas aeruginosa chitinase, a gradually secreted protein. J Bacteriol, 2001. 183(24): p. 7044–52.

60. Deziel, E., et al., Analysis of Pseudomonas aeruginosa 4-hydroxy-2-alkylquinolines (HAQs) reveals a role for 4-hydroxy-2-heptylquinoline in cell-to-cell communication. Proc Natl Acad Sci U S A, 2004. 101(5): p. 1339–44.

61. Calfee, M.W., J.P. Coleman, and E.C. Pesci, Interference with Pseudomonas quinolone signal synthesis inhibits virulence factor expression by Pseudomonas aeruginosa. Proc Natl Acad Sci U S A, 2001. 98(20): p. 11633–7.

62. Moradali, M.F., S. Ghods, and B.H. Rehm, Pseudomonas aeruginosa Lifestyle: A Paradigm for Adaptation, Survival, and Persistence. Front Cell Infect Microbiol, 2017. 7: p. 39.

63. Venkataraman, A., et al., Metabolite transfer with the fermentation product 2,3-butanediol enhances virulence by Pseudomonas aeruginosa. ISME J, 2014. 8(6): p. 1210–20.

64. Blair, J.M., et al., Expression of homologous RND efflux pump genes is dependent upon AcrB expression: implications for efflux and virulence inhibitor design. J Antimicrob Chemother, 2015. 70(2): p. 424–31.

65. Wang, J.D. and P.A. Levin, Metabolism, cell growth and the bacterial cell cycle. Nat Rev Microbiol, 2009. 7(11): p. 822–7.

66. Lamarche, M.G. and E. Deziel, MexEF-OprN efflux pump exports the Pseudomonas quinolone signal (PQS) precursor HHQ (4-hydroxy-2-heptylquinoline). PLoS ONE, 2011. 6(9): p. e24310.

67. Farrow, J.M., 3rd and E.C. Pesci, Two distinct pathways supply anthranilate as a precursor of the Pseudomonas quinolone signal. J Bacteriol, 2007. 189(9): p. 3425–33.

68. Raychaudhuri, A., A. Jerga, and P.A. Tipton, Chemical mechanism and substrate specificity of RhlI, an acylhomoserine lactone synthase from Pseudomonas aeruginosa. Biochemistry, 2005. 44(8): p. 2974–81.

69. Diaz-Perez, A.L., et al., The gnyRDBHAL cluster is involved in acyclic isoprenoid degradation in Pseudomonas aeruginosa. Appl Environ Microbiol, 2004. 70(9): p. 5102–10.

70. Kanehisa, M., et al., New approach for understanding genome variations in KEGG. Nucleic Acids Res, 2019. 47(D1): p. D590–D595.

71. Kang, Y., et al., The long-chain fatty acid sensor, PsrA, modulates the expression of rpoS and the type III secretion exsCEBA operon in Pseudomonas aeruginosa. Mol Microbiol, 2009. 73(1): p. 120–36.

72. Oshri, R.D., et al., Selection for increased quorum-sensing cooperation in Pseudomonas aeruginosa through the shut-down of a drug resistance pump. ISME J, 2018.

73. Tian, Z.X., et al., Transcriptome profiling defines a novel regulon modulated by the LysR-type transcriptional regulator MexT in Pseudomonas aeruginosa. Nucleic Acids Res, 2009. 37(22): p. 7546–59.

74. Simon, R., et al., Plasmid vectors for the genetic analysis and manipulation of rhizobia and other gram-negative bacteria. Methods Enzymol, 1986. 118: p. 640–59.

75. Kirke, D.F., et al., The Aeromonas hydrophila LuxR homologue AhyR regulates the N-acyl homoserine lactone synthase, AhyI positively and negatively in a growth phase-dependent manner. FEMS Microbiol Lett, 2004. 241(1): p. 109–17.

76. Swift, S., et al., Quorum sensing in Aeromonas hydrophila and Aeromonas salmonicida: identification of the LuxRI homologs AhyRI and AsaRI and their cognate N-acylhomoserine lactone signal molecules. J Bacteriol, 1997. 179(17): p. 5271–81.

77. Linares, J.F., et al., Overexpression of the multidrug efflux pumps MexCD-OprJ and MexEF-OprN is associated with a reduction of type III secretion in Pseudomonas aeruginosa. J Bacteriol, 2005. 187(4): p. 1384–91.

78. Fletcher, M.P., et al., A dual biosensor for 2-alkyl-4-quinolone quorum-sensing signal molecules. Environ Microbiol, 2007. 9(11): p. 2683–93.

79. Becher, A. and H.P. Schweizer, Integration-proficient Pseudomonas aeruginosa vectors for isolation of single-copy chromosomal lacZ and lux gene fusions. Biotechniques, 2000. 29(5): p. 948–50, 952.

80. Gould, T.A., H.P. Schweizer, and M.E. Churchill, Structure of the Pseudomonas aeruginosa acyl-homoserinelactone synthase LasI. Mol Microbiol, 2004. 53(4): p. 1135–46.

